# Beyond the Resection: Surgical White Matter Disruption Structurally Alters Non-Resected Brain Anatomy

**DOI:** 10.1101/2025.03.19.643850

**Authors:** Philip Pruckner, Remika Mito, David Vaughan, Graeme Jackson, Florian Fischmeister, Karl Heinz-Nenning, Marc Berger, Ekaterina Pataraia, Christoph Baumgartner, Christian Dorfer, Karl Rössler, Thomas Czech, Gregor Kasprian, Silvia Bonelli, Robert Smith

## Abstract

Resective neurosurgery is a cornerstone treatment for many neurological conditions. Although traditionally viewed as localised procedure, increasing evidence from advanced magnetic resonance imaging (MRI) shows that also non-resected anatomy can degenerate following surgery. The relationship between local tissue removal and these postoperative changes remains thus far speculative. Here, we investigate the hypothesis that degenerative changes to surgically preserved grey and white matter are mediated by *transneuronal degeneration*, a deterioration of intact neuronal populations due to lost axonal input. Using a robust structural and diffusion MRI framework, we first identify widespread postoperative atrophy: pronounced cortical thickness decreases near the resection, and extensive white matter impairments across the ipsilateral hemisphere. Importantly, we then link these alterations to surgical white matter disruption, revealing a sequential network atrophy following neurosurgery. Beyond degenerative effects, we also demonstrate often reported structural network reorganisations as an artefact of image processing, indicating limited capacity for macroscale plasticity post-resection.

**Key Points:**

- Resective neurosurgery can lead to postoperative atrophy of non-resected grey and white matter. Utilising a robust longitudinal neuroimaging framework, we evaluate these changes as consequences of *transneuronal degeneration*, demonstrating a sequential atrophy of brain anatomy following the loss of surgically disrupted white matter.
- The majority of severed connections projected to atrophic cortices near the resection, indicating them to be short association fibers. Consecutive white matter atrophy however affected extensive ipsilateral pathways, highlighting a cascading network effect of resective brain surgery.
- We did not find evidence to support often reported structural reorganisations of large-scale brain networks post-resection. Yet, we demonstrate that evidence typically interpreted as such can be replicated when using non-robust reconstruction methods.

## Main

Each year, over 13 million neurosurgical procedures are required worldwide, making brain surgery a fundamental treatment for many neurological conditions^1^. A substantial proportion of these cases – spanning tumors, vascular malformations or epilepsy – necessitate the resection of diseased structures. This demands a careful balance: removing sufficient pathological tissue to treat underlying conditions while preserving critical brain function. Traditionally this is achieved by sparing eloquent structures from resection; however, emerging evidence from advanced magnetic resonance imaging (MRI) suggests that even non-resected anatomy can degenerate, documenting postoperative grey matter atrophy^2–5^, along with extensive white matter impairments^6–8^. The relationship between local tissue resection and these postoperative changes remains poorly characterised.

One possible explanation for anatomical alterations outside the immediate site of resection may be the surgical disruption of axonal pathways^3^. Axons are elongated projections of neurons to distant brain regions, serving as conduits for electrical signalling as well as for the transport of trophic and metabolic factors. When these connections *(or their respective cell bodies)* are injured during resection, they degenerate^9^. This results in a loss of electrical stimulation and diminished nutritional support for disconnected neurons, ultimately leading to a postoperative deterioration of otherwise intact cell populations. This process is known as *transneuronal degeneration*^10^ and has been proposed as one of the key mechanisms facilitating the propagation of pathologies along brain networks^11^ *(for a schematic, see Fig. 1)*.

**Fig. 1.**
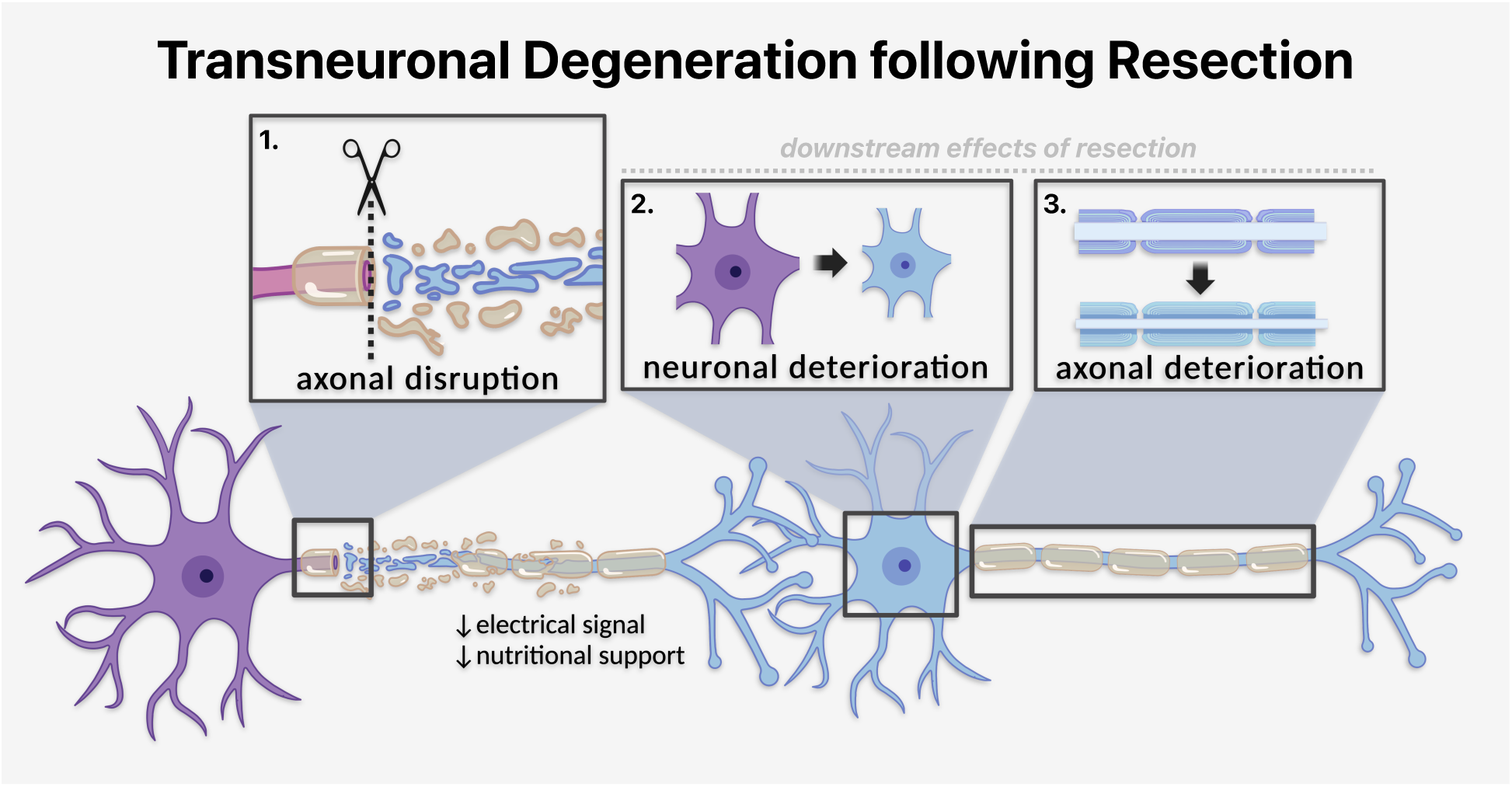
Transneuronal degeneration following surgical tissue resection. Illustrated is the proposed mechanism driving atrophy along postoperatively preserved grey and white matter: 1) the surgical disruption of axons leads to their degeneration. This results in loss of electrical signal and nutritional support for neurons downstream of resection, ultimately leading to a 2) deterioration of otherwise intact neurons and 3) axonal projections thereof.

Despite the mechanism’s conceptual appeal to explain structural change beyond the resection, it has been difficult to assess in neurosurgical studies. Firstly, the cortical projections of resected white matter are clinically unknown, as they are neither apparent during intervention nor assessed postoperatively. Scientifically they are likely under-reported, as previous research has predominately focused on injury of major bundles^12,13^, which are only a subset of the brain’s extensive neural networks^14^. It therefore remains unclear if anatomical alterations align with endpoints of resected connective tissue. Moreover, evidence of transneuronal degeneration primarily stems from animal studies^10^; whether it applies across the brain’s ubiquitous neuronal pathways is yet to be determined, as robust human evidence is restricted to studies of the visual system^15–18^.

In this article, we use advanced structural and diffusion MRI methods to evaluate postoperative neuroanatomical alterations as consequences of surgically resected white matter. Specifically, we investigate structural changes following two common resective treatments for temporal lobe epilepsy: transsylvian selective amygdalohippocampectomy^19–21^ and anterior temporal lobectomy^22^. Epilepsy surgery serves as a particularly well-suited context for such inquiry, as compared to other indications for resective surgery, subjects typically present without gross morphological abnormalities that would confound analysis. Utilising a tailored longitudinal imaging framework, we estimate three structural effects that we hypothesise to sequentially manifest following local tissue resection, namely: *(1)* the loss of surgically disrupted white matter *(primary effect), (2)* the postoperative atrophy of non-resected cortices *(secondary effect)* as well as *(3)* the postoperative atrophy of non-resected white matter *(tertiary effect).* Through these structural effects we posit a biologically motivated imaging model of transneuronal degeneration, quantitatively addressing the central question: can surgical disruption of axonal pathways explain grey and white matter alterations downstream of resection?

## Results

### Sample characteristics and experimental overview

We investigated 59 patients who underwent resective brain surgery for temporal lobe epilepsy at the Medical University of Vienna between 2012 and 2019. Patients were treated with either transsylvian selective amygdalohippocampectomy^19–21^ *([ts]SAHE, left:right=16:12)* or anterior temporal lobectomy^22^ *(ATL, left:right=11:20).* Surgery type was determined by an interdisciplinary epilepsy board based on individual clinical, electrophysiological, and imaging findings. Baseline characteristics did not differ between groups, except that more male patients underwent ATL *(see Methods)*.

Preoperative T1-weighted *(T1w)* and diffusion-weighted MRI *(dMRI)* were used to estimate the individual white matter lost to resection *(primary effect).* Longitudinal cortical thickness changes *(secondary effect)* were assessed with postoperative T1w MRI, and longitudinal connectivity changes in non-resected white matter *(tertiary effect)* were assessed with postoperative dMRI. The requisite imaging data to assess primary/secondary/tertiary effects was available in 59 *(ATL:SAHE=31:28)*, 51 *(ATL:SAHE=28:23)* and 35 participants *(ATL:SAHE=15:20),* respectively.

Each of these effects was estimated for non-resected parts of brain regions, as defined by the Desikan-Killiany^23^ parcellation, along with subcortical parcels where applicable. Results were aggregated across subjects of each treatment group based on the laterality of resection, differentiating between ipsi- and contralateral hemispheres. The relationships between estimated effects were explored within an imaging model of transneuronal degeneration, testing the explanatory power of white matter lost to resection *(primary effect)* for postoperative changes *(secondary and tertiary effects, see Fig. 2A)*.

**Figure 2.**
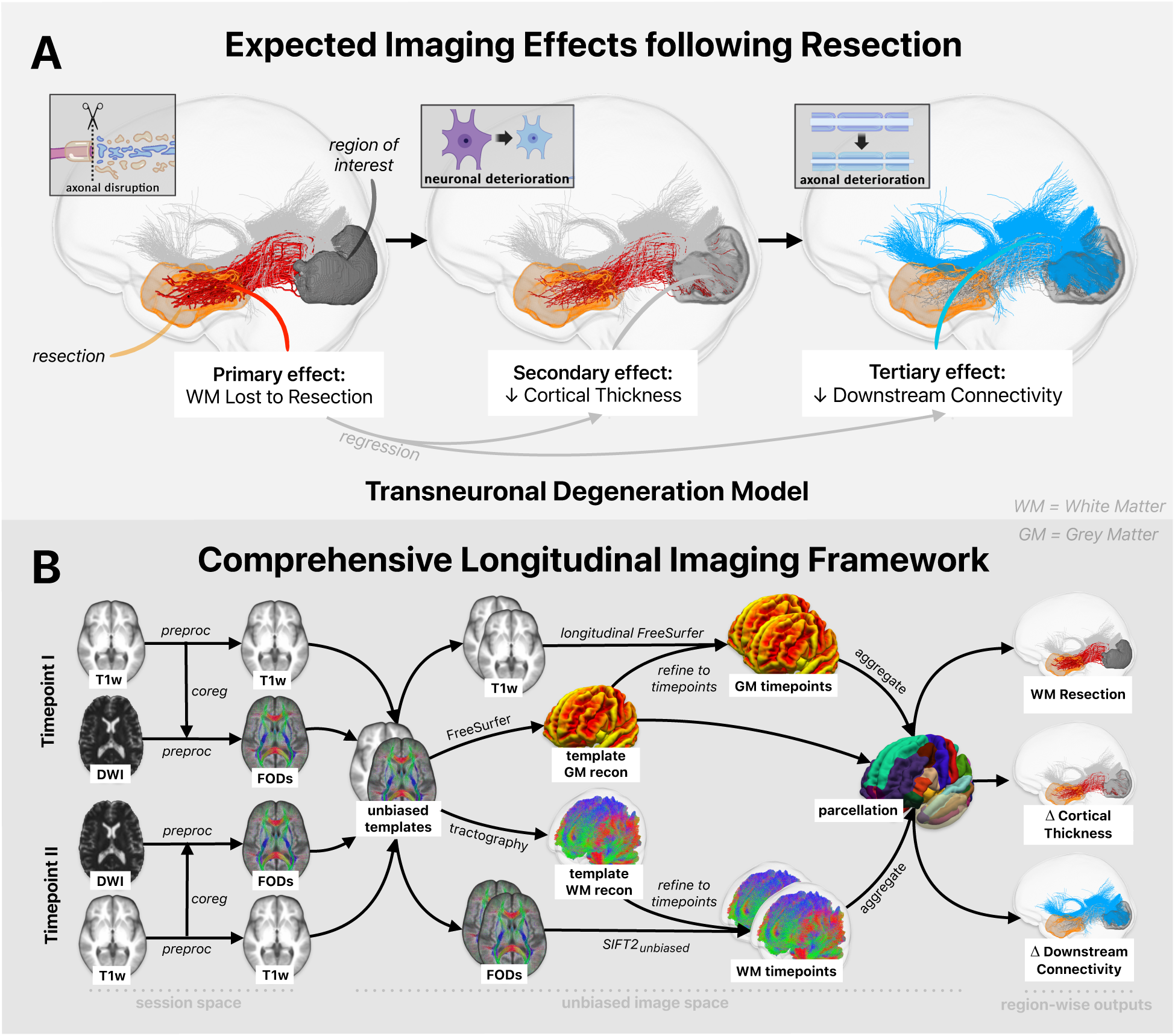
Expected imaging effects following tissue resection and employed longitudinal analysis framework. A) Expected imaging effects following tissue resection for an exemplar, non-resected brain region. The primary effect is the loss of surgically resected white matter, leading to degeneration of affected connections. This induces secondary and tertiary effects, downstream cortical atrophy due to a degeneration of disconnected neurons and downstream white matter atrophy due to a degeneration of axonal projections thereof. These effects were used to construct an imaging model of transneuronal degeneration, testing the explanatory power of white matter lost to resection for downstream effects. B) Comprehensive imaging framework for robust longitudinal analysis of grey and white matter, integrating an adapted longitudinal Freesurfer and the novel SIFT2_unbiased_ framework: preoperative and postoperative data were preprocessed and spatially transformed to unbiased subject templates. Relevant grey and white matter properties were reconstructed on these templates and refined to fit individual session data; this allows reliable estimation of longitudinal features across modalities while maintaining spatial correspondence. Note that additional steps were taken to adjust for the complexities of surgical image data. Abbreviations: DWI = diffusion-weighted-imaging, FOD = fiber orientation distribution, GM = grey matter, WM = white matter, postop = postoperative, preop = preoperative, recon = reconstruction, SIFT = spherical deconvolution-based filtering of tractograms.

For reliable estimation of expected grey and white matter effects, we utilised a robust longitudinal imaging framework that integrates novel connectivity reconstruction methods with adapted cortical analysis tools; an overview of the employed imaging pipeline is shown in *Fig. 2B*. Note that additional steps were taken to address the complexities of gross resections in surgical image data. Further clinical details as well as technical descriptions of the methods used for this study are provided in the *Methods* and *Supplementary Materials*.

### Local tissue resections and definition of surgically preserved regions

The goal of temporal lobe epilepsy surgery is to remove structures involved in epileptogenesis, typically targeting the amygdala, hippocampus as well as adjacent cortices. While ATL is performed as extensive *en-bloc* resection of the anterior temporal lobe, tsSAHE removes target structures through a limited resection via the Sylvian fissure. To define the extent of tissue resections, we segmented each subject’s resection zone from spatially aligned pre- and postoperative T1w images, quantifying the intersection of preoperative brain tissue with the *(cerebrospinal fluid filled)* postoperative resection cavity *(see Supplementary Materials)*. All generated resection masks were inspected and manually corrected if necessary.

To visualise the spatial distribution of tissue resections, we aggregated all resection masks within a common template *(fsaverage),* plotting the tissue removed in ≥ 10% of subjects for each treatment group. Penetrance maps illustrate the corresponding frequencies of tissue resection. As clinically intended, ATL removed more tissue than tsSAHE *(Fig. 3A)*.

**Figure 3.**
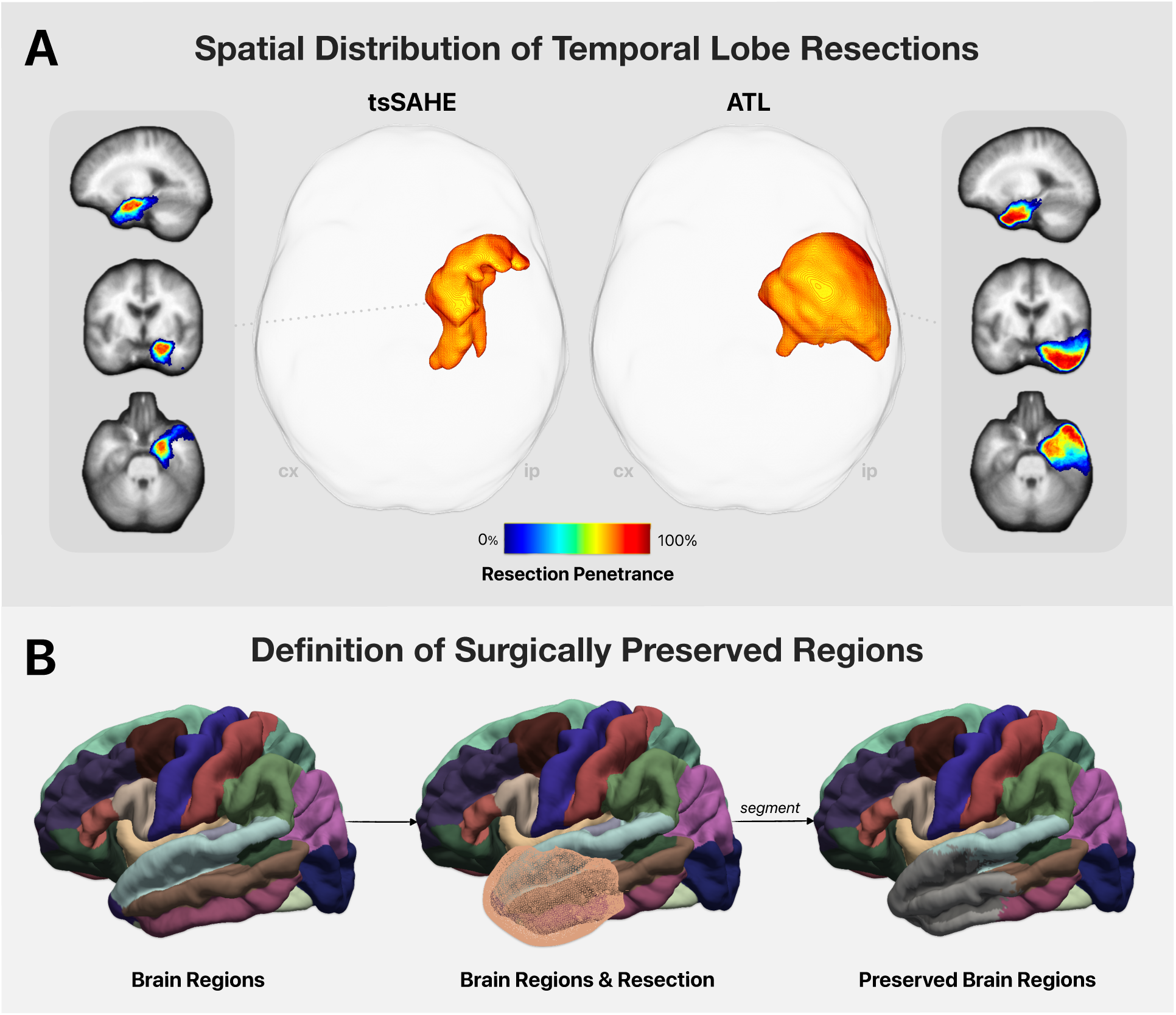
Volumetric tissue resection and definition of surgically preserved regions. A) Individual resection cavity segmentations were aggregated within a common template. Axial glass brain views show the volumetric tissue removed in at least l0% of participants. Penetrance maps on the sides show the corresponding resection frequencies. As clinically intended, more temporal lobe tissue was removed after ATL. B) For each participant we bisected anatomical brain regions into resected and surgically preserved parts based on their intersections with the resection cavity. ATL = anterior temporal lobectomy, cx = contralateral, ip = ipsilateral, tsSAHE = transsylvian selective amygdalohippocampectomy.

Using these resection masks, we next bisected each subject’s brain regions into resected and non-resected parts *(Fig. 3B)*. All subsequent estimations of structural effects that we hypothesised to arise following tissue resection *(primary, secondary and tertiary effects)* were exclusively performed for the non-resected portion of brain regions. These non-resected parts of brain regions will be from here on referred to as *surgically preserved*.

### Primary Effect: Regional White Matter Lost to Resection

To demonstrate the trajectories of surgically disrupted white matter fibers, we reconstructed a streamline tractogram from cohort-averaged fibre orientation distributions and determined the set of streamlines that intersected the resection cavity in ≥ 10% of subjects; streamlines were color-coded according to their resection penetrance *(Figure 4A).* Despite their distinctly different tissue resections, ATL and tsSAHE both led to extensive white matter disruptions, mostly affecting bundles that connected regions within the ipsilateral hemisphere.

**Figure 4.**
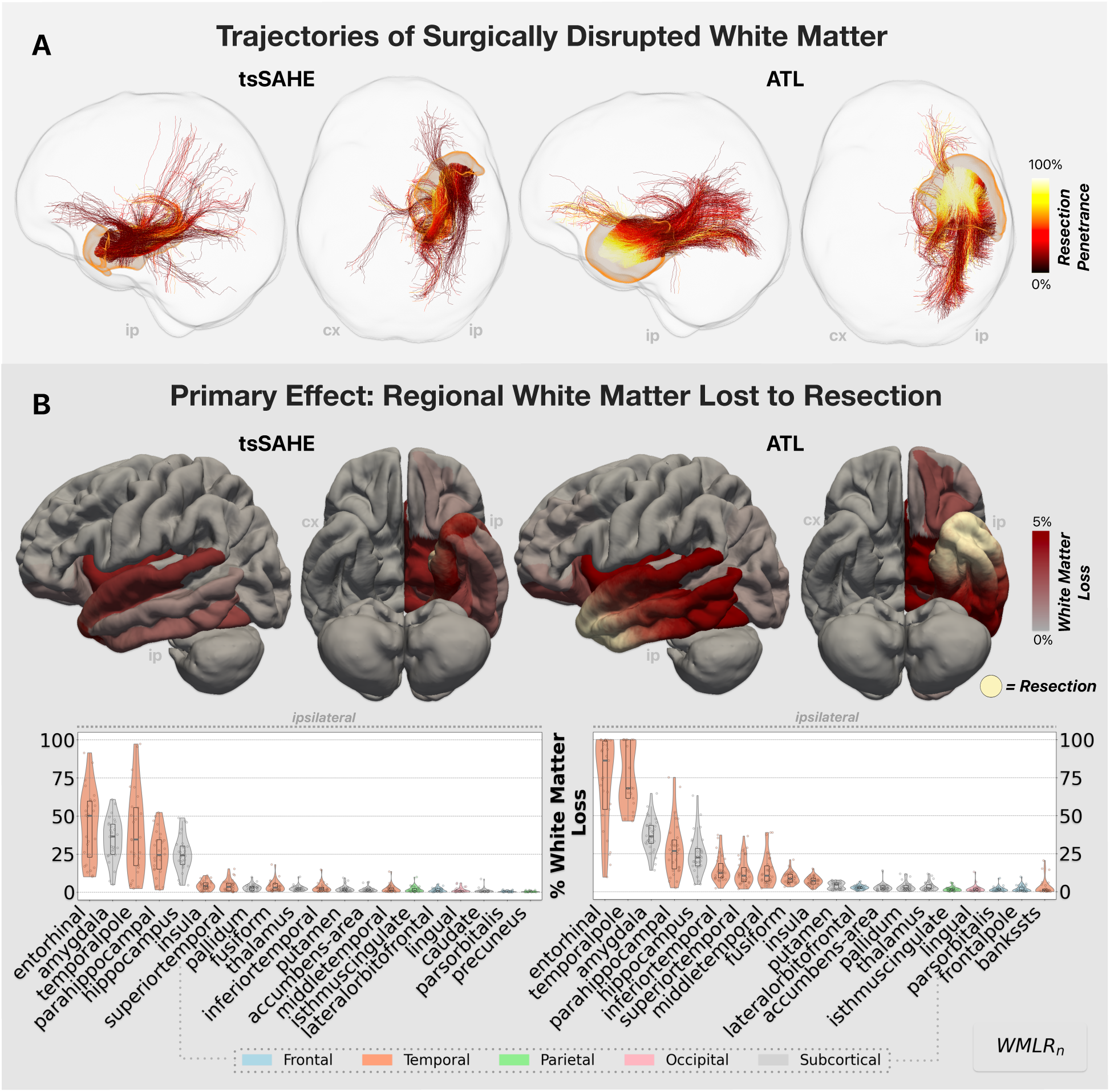
Primary Effect: Regional White Matter Lost to Resection. A) Estimated trajectories of white matter bundles disrupted during resection, visualised using streamline tractography. A whole brain tractogram was constructed from cohort-averaged fibre orientation data; streamlines intersecting resection cavities of ≥ 10% of subjects are visualised and coloured according to their resection penetrance. Despite their different resection volumes, both treatments led to extensive white matter injuries. B) Brain surfaces: regional White Matter Lost to Resection (primary effect, *WMLR*_*n*_) plotted for each treatment group. Vertices within the (treatment group average) resection cavity were additionally weighted according to their resection penetrance. Violin plots: Plotted are the top 20 grey matter regions for each surgical technique, with x-axes sorted by the median white matter losses. Violins are cropped at the minimal and maximal values. Altogether, we found greatest white matter losses for regions close to resection, indicating the majority of severed bundles to be short association fibers. Abbreviations: ATL = anterior temporal lobectomy, cx = contralateral, GM = grey matter, ip = ipsilateral, tsSAHE = transsylvian amygdalohippocampectomy, WMLR_n_ = regional White Matter Lost to Resection.

To quantitatively assess the cortical projections of surgically disrupted white matter, we evaluated the regional White Matter Lost to Resection *(primary effect):* for each preserved cortical and subcortical region *n*, we identified streamlines within individual *(SIFT2 weighted*^24^) preoperative tractograms that intersected the resection cavity, determining the fraction of Fibre Bundle Capacity *(a measure of intra-axonal cross-sectional area*^25^) involved in resection. This quantitative measurement is from here on referred to as regional White Matter Lost to Resection *(WMLR*_*n*_, *see Figure 4B)*. We found extensive losses of white matter following both surgical treatments, with greatest effects sizes observed for regions close to tissue resection. For tsSAHE, white matter losses of ipsilateral temporal lobe regions ranged between 1% *(middle temporal cortex)* and 50% *(entorhinal cortex)*, whereas for ATL white matter losses ranged between 7% *(insula)* and 86% *(entorhinal cortex)*. Extratemporal white matter losses were <3% after tsSAHE and <5% following ATL.

These results highlight that local tissue resection can lead to pronounced disruptions of axonal pathways. Especially regions close to the site of resection experienced pronounced losses of white matter, suggesting that the majority of disrupted bundles were short association fibers.

### Secondary Effect: Regional Cortical Thickness Change

Next, we evaluated the presence of postoperative cortical atrophy. We adapted the longitudinal Freesurfer^26–28^ framework for image data containing gross resection *(see Methods)* and estimated the pre- and postoperative cortical thickness for grey matter regions. For each preserved cortical region *n*, we calculated the fractional change in Cortical Thickness *(secondary effect,* Δ*CT*_*n*_) and tested for statistical longitudinal differences *(paired t-test with FDR correction, see Fig. 5)*.

**Figure 5.**
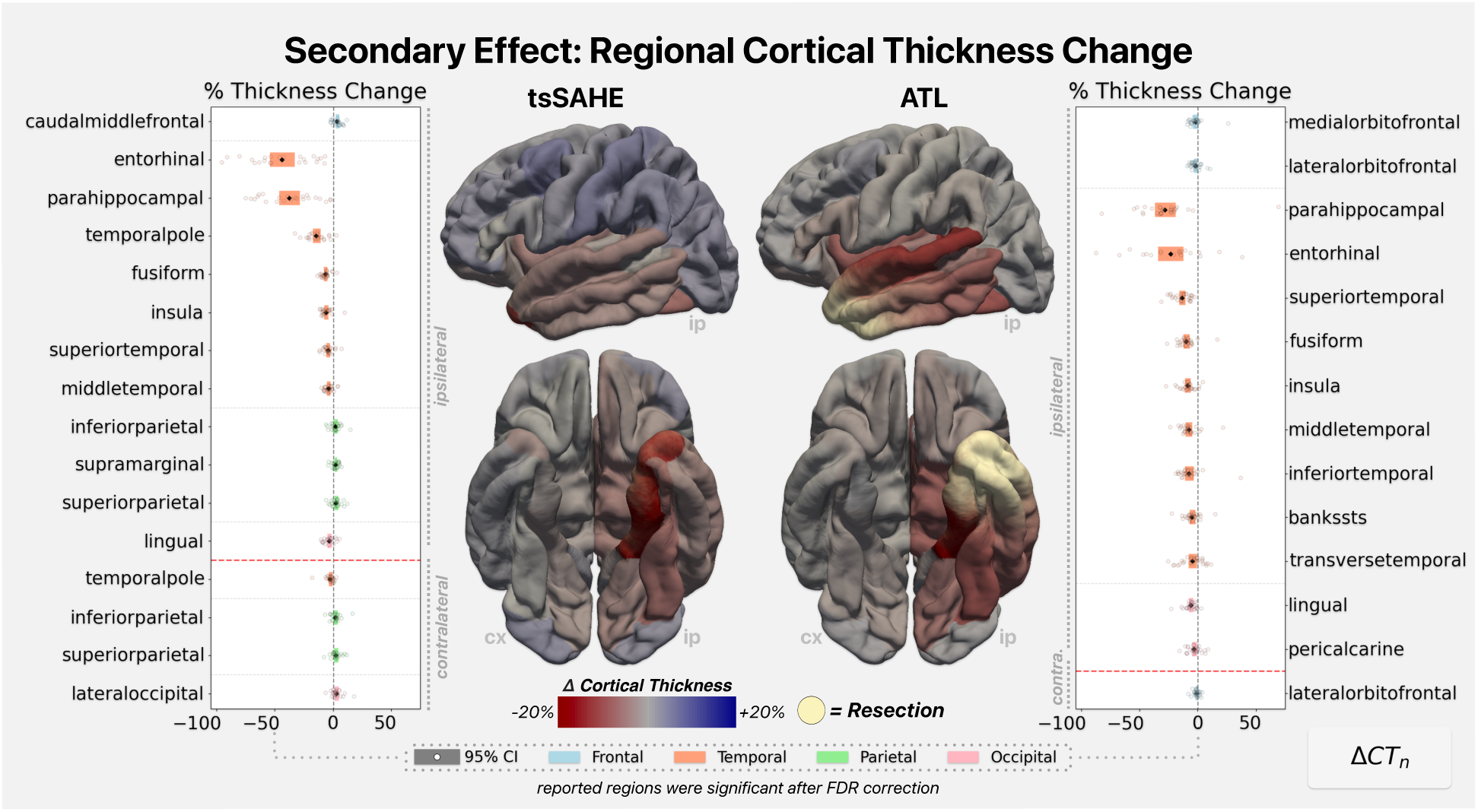
Secondary Effect: Regional Cortical Thickness Change. Longitudinal change in cortical thickness (Δ*CT*_*n*_), summarised per brain region for each treatment group. Bar plots: mean and 95% confidence intervals of regions with significant longitudinal change (paired t-test, FDR corrected), categorised by brain lobes. Brain surfaces: mean cortical thickness change summarised per brain region for each treatment group. Vertices within the (treatment group average) resection cavity were additionally weighted according to their resection penetrance. We found pronounced postoperative atrophy, especially for regions close to the resection cavity. Abbreviations: ATL = anterior temporal lobectomy, cx = contralateral, ip = ipsilateral, tsSAHE = transsylvian selective amygdalohippocampectomy.

We identified significant cortical atrophy across both treatment groups, which was, again, most pronounced in the vicinity of resection. For tsSAHE, the majority of ipsilateral temporal lobe regions showed significant cortical thinning, with effect sizes between 4% *(middle temporal cortex)* and 44% *(entorhinal cortex).* After ATL, all except for one ipsilateral temporal lobe region showed significant cortical thickness decreases, with effect sizes between 5% *(transverse temporal cortex)* and 28% *(parahippocampal cortex).* Effect sizes within ipsilateral extratemporal cortices were <4% for tsSAHE and <7% for ATL, respectively. Despite the expected postoperative atrophy, a number of significant regional cortical thickness increases were also found after tsSAHE, with effect sizes <3%.

Altogether, longitudinal analysis of cortical thickness revealed significant postoperative atrophy that spatially matched the estimated projections of severed white matter bundles. A few regions exhibited a slight increase in cortical thickness after targeted surgery, although interpretability of this finding is limited in the absence of a clear *a-priori* hypothesis.

### Transneuronal degeneration model: loss of axonal input is related to cortical atrophy

Thus far, we demonstrated comparable patterns of postoperative cortical atrophy and projections of surgically resected white matter. To statistically evaluate the relationship between *WMLR_n_ (primary effect)* and Δ*CT_n_ (secondary effect)* we used a mixed linear effect model with random intercepts, testing for a cohort-wide effect across 2748 brain regions of 51 patients that had streamline intersections with the resection cavity (*ATL:SAHE=28:23)*. *WMLR_n_ (log_10_-transformed to ensure model linearity*) and clinical covariates, were included as fixed effects. Subjects were included as random effect to account for the hierarchical data structure *(see Methods)*.

We found *log*_10_*WMLR_n_ (primary effect)* to be significantly correlated with Δ*CT*_*n*_ *(secondary effect)*, with every 10-fold loss of resected white matter corresponding to a 3.4% decrease in cortical thickness *(p<0.0001; see Figure 7A).* We found no association between clinical covariates and cortical thickness change. Fixed effects explained 19% *(R^2^_marginal_)* and random effects 9% of variance. Altogether, fixed and random effects explained 28% of variance *(R^2^_conditional_)*.

The analysis supports the idea that the surgical disruption of white matter connections leads to postoperative atrophy in cortices downstream of resection.

### Tertiary Effect: Downstream Connectivity Decline

We further evaluated the presence of postoperative atrophy within the non-resected white matter. To measure this, we generated pre- and postoperative quantitative *(SIFT2 weighted*^24^) tractograms using a robust longitudinal diffusion MRI framework specifically designed for this study *(SIFT2_unbiased_, see Methods)*. For each preserved cortical and subcortical region *n*, we determined the total pre- and postoperative Fibre Bundle Capacity of streamlines downstream of resection – that is, specifically those not intersecting the resection zone. Then we calculated the fractional change in Downstream Connectivity (Δ*DC*_!,_ *tertiary effect)* and tested for statistical longitudinal differences *(paired t-test with FDR correction, see Figure 6)*.

**Figure 6.**
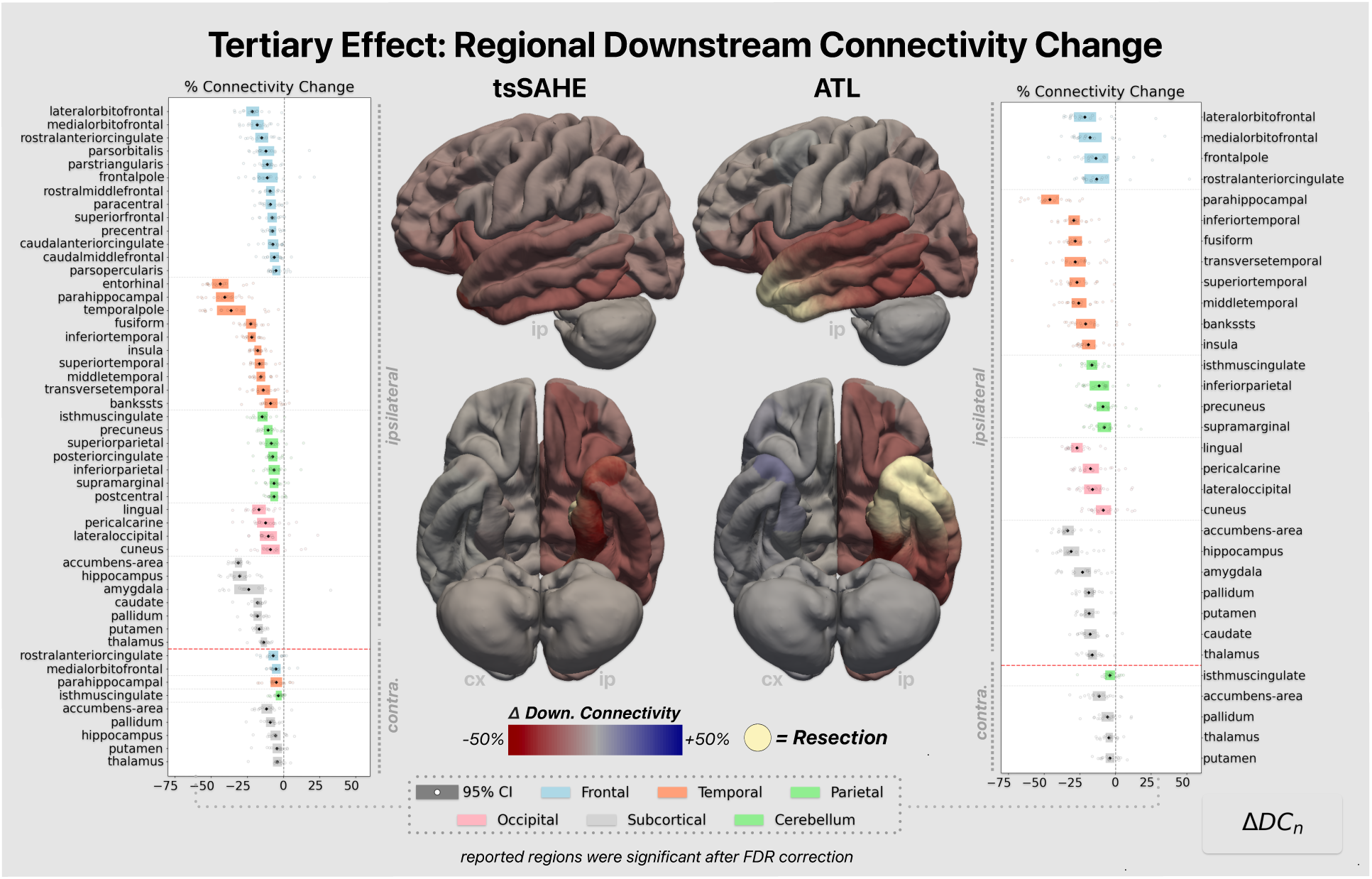
Tertiary effect: Downstream connectivity change. Longitudinal connectivity change in surgically preserved white matter (Δ*DC_n_*), summarised per brain region for each treatment group. Bar plots: mean and 95% confidence intervals of regions with significant longitudinal change (paired t-test, FDR corrected), categorised by brain lobes. Brain Surfaces: mean downstream connectivity change summarised per brain region for each treatment group. Vertices within the (treatment group average) resection cavity were additionally weighted according to their resection penetrance. We found extensive postoperative white matter atrophy mainly within the hemisphere ipsilateral to resection. Abbreviations: ATL = anterior temporal lobectomy, cx = contralateral, ip = ipsilateral, tsSAHE = transsylvian selective amygdalohippocampectomy.

We found widespread significant connectivity changes - exclusively decreases – for regions of both treatment groups, mainly located within the ipsilateral hemisphere but also extending contralaterally. For tsSAHE, all ipsilateral brain regions exhibited significantly decreased downstream connectivity, for ATL the majority did. While strongest effects were again most notable within the ipsilateral temporal lobes, with up to 46% for tsSAHE *(entorhinal cortex)* and ATL *(parahippocampal cortex),* now also a high number of extratemporal ipsilateral regions showed strong effect sizes of up to 34%.

Our longitudinal analysis of downstream connectivity revealed extensive atrophy of surgically preserved axonal pathways. Compared to *primary* and *secondary* effects, we found an increased number of significantly altered regions as well as increased effect sizes, indicating a cascading effect of resective brain surgery on the white matter not involved in resection. No significant connectivity increases were identified, suggesting that previously described hypertrophic cortices were not part of larger network reorganisations.

### Transneuronal degeneration model: loss of axonal input is related to downstream connectivity change

Finally, we evaluated if the surgical disruption of axonal pathways is associated with postoperative atrophy of surgically preserved white matter. Using mixed effects model with random intercepts, we tested for a cohort-wide effect between *WMLR_n_* (*primary effect)* and Δ*DC_n_ (tertiary effect)* across 2409 brain regions of 35 patients that had streamline intersections with the resection cavity *(ATL:SAHE=15:20).* As before, *WMLR_n_ (log_10_-transformed to ensure model linearity)* and clinical covariates were included as fixed effects and subjects as random effect *(see Methods)*.

We found *log*_10_ *WMLR_n_ (primary effect)* to significantly correlate with Δ*DC_n_ (tertiary effect),* with every 10-fold loss of white matter to resection corresponding to a 7.2% connectivity decrease in white matter downstream of resection *(p<0.0001, see Fig. 7B).* We found no association between clinical covariates and downstream connectivity change. Fixed effects were able to explain 34% of variance *(R^2^_marginal_)*, random effects 24% of variance. Altogether, fixed and random effects explained 58% of variance *(R^2^_conditional_)*.

**Figure 7.**
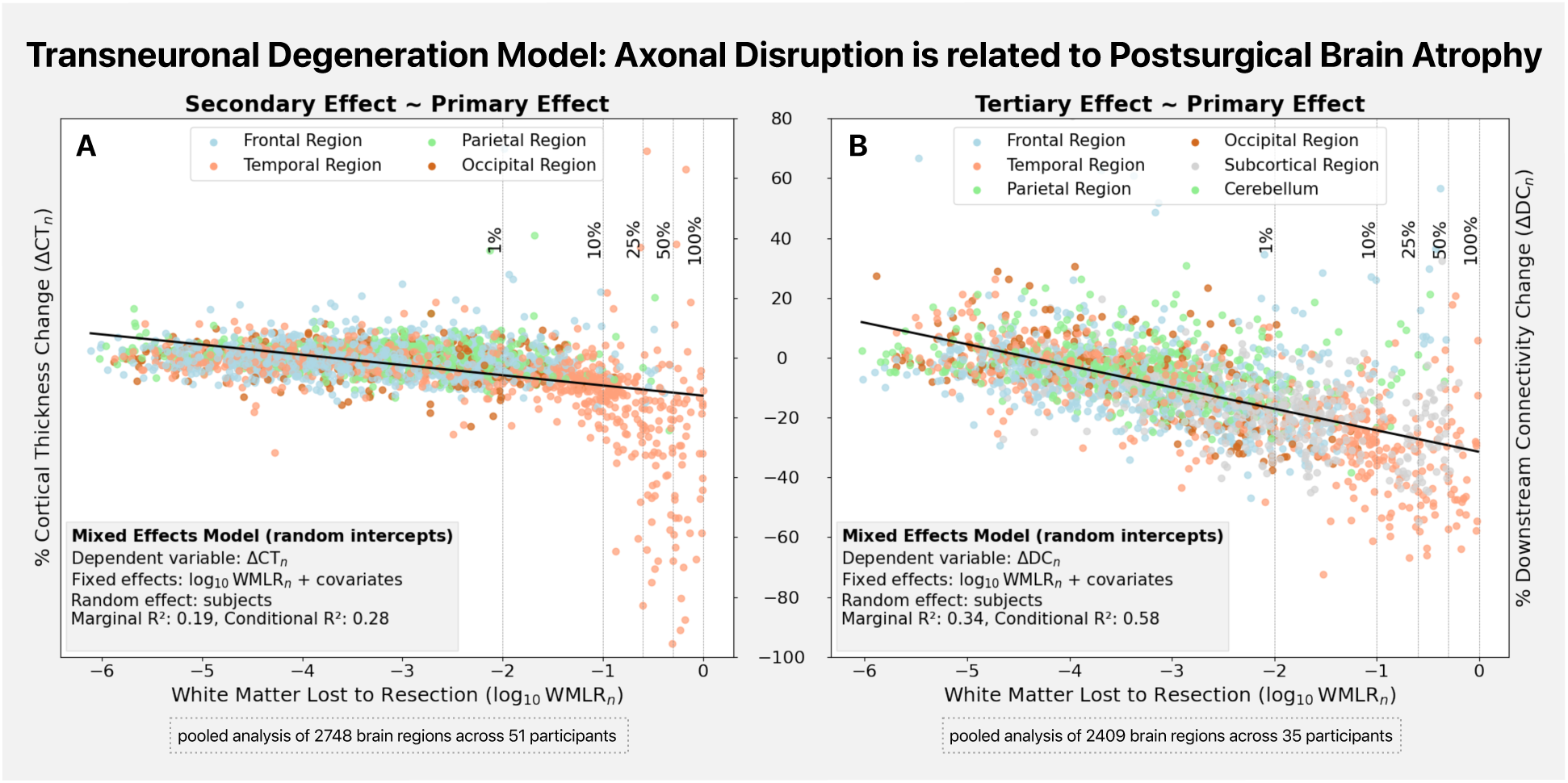
Imaging model of transneuronal degeneration. Mixed effects models exploring the relationships between White Matter Lost to Resection (*log*_10_ *WMLR*_n_) and change in Cortical Thickness (Δ*CT_n_*) as well as Downstream Connectivity (Δ*DC_n_*). Regional white matter losses and clinical covariates were included as fixed effects, subjects were included as random effect. A) Relationship between resected white matter and longitudinal decreases in cortical thickness. B) Relationship between resected white matter and longitudinal decreases in downstream connectivity. Surgical resection of white matter was closely related to grey and white matter atrophy in brain regions downstream of resection.

The result shows that surgical white matter disruption has an impact on the postoperatively preserved white matter, extending the structural effect of resection beyond primarily disconnected synapses.

## Discussion

The results presented in this article emphasise the brain-wide consequences of resective neurosurgery, identifying the surgical injury of white matter as a key factor driving structural change beyond the immediate resection. Across two surgical interventions and image modalities, we demonstrated that surgical resection of axonal pathways is consistently associated with postoperative atrophy in non-resected cortices and impairment of downstream white matter, indicating a systematic degeneration of anatomical networks that extends beyond primarily disconnected synapses. Beyond degenerative effects, our analysis did not reveal evidence for large-scale structural reorganisations of white matter networks, challenging previous reports of such phenomena. The novel longitudinal brain connectivity tools developed for this study integrated seamlessly with existing structural pipelines, providing a powerful framework to study neuroanatomical change.

### The structural effect of resection propagates throughout the brain

Non-resected neuroanatomy can degenerate after surgery^2–8^, which we hypothesised to occur as a consequence of *transneuronal degeneration*, where intact neurons deteriorate due to the loss of axonal input. Using advanced neuroimaging methods, we measured the expected imaging manifestations of this mechanism, which were then used to construct a quantitative model that tests its presence following two major types of resective epilepsy surgery, anterior temporal lobectomy *(ATL)* and transsylvian selective amygdalohippocampectomy *(tsSAHE)*.

First, we identified the extent of severed white matter following each procedure, demonstrating widespread disruptions of intrahemispheric connections, even after restricted tissue resection. While visual inspection of streamlines that intersected the resection cavity emphasised the involvement of long-distance bundles *(see Fig. 4A),* quantitative streamline tractography^25^ revealed that the majority of severed bundles projected to regions close to resection *(primary effect, see Fig. 4B)*. This suggests that short association fibers – estimated to account for 60% of the brain’s white matter^14^ – are the predominant fiber type injured during tissue resection. The finding highlights the extensive white matter disruptions that arise from resective surgery, and most importantly, challenges the prevailing clinical and research focus on major long-distance tracts.

As others have, we demonstrated significant postoperative atrophy of non-resected grey matter^2–5^ *(secondary effect, see Fig. 5)*. Importantly though, we also showed spatial correspondence of cortical atrophy with projections of severed white matter *(for a visual comparison, see Fig. 4 and 5).* Regression analysis statistically confirmed this relationship, showing a close association between the regional White Matter Lost to Resection *(primary effect)* and the regional change in Cortical Thickness *(secondary effect, see Fig. 7A)*.

While the overall relationship was well approximated by our log-linear model, regions with white matter losses of >10% experienced accelerated cortical thinning; in-fact, re-fitting the same model exclusively for these regions found cortical thinning to be increased by a factor of 6.2 *(Extended Data Fig. 1)*. This suggests a potential vulnerable threshold, above which cortices experience more pronounced cortical atrophy.

Moreover, we also found extensive postoperative impairments of non-resected white matter^6–8^ *(tertiary effect, see Figure 6)*, which predominately affected connections across the ipsilateral hemisphere. The regional decrease of Downstream Connectivity *(tertiary effect)* was associated with the regional White Matter Lost to Resection *(primary effect, see Figure 7B)*, indicating the structural effect of resection to extend into the surgically preserved white matter. The spatial extent and effect sizes of regional changes in Downstream Connectivity *(tertiary effect)* exceeded those of previously described effects, suggesting a cascading impact on non-resected pathways.

Together, we show extensive grey and white matter atrophy after resective surgery, even following restricted resections. Our imaging model attributed these changes to the surgical resection of white matter, indicating atrophy to sequentially propagate along the brain’s interconnected anatomical network following the loss of axonal input. These findings align closely with the mechanism of *transneuronal degeneration*^10,11^, which we propose to drive degenerative changes post-surgery.

**Extended Data Fig 1.**
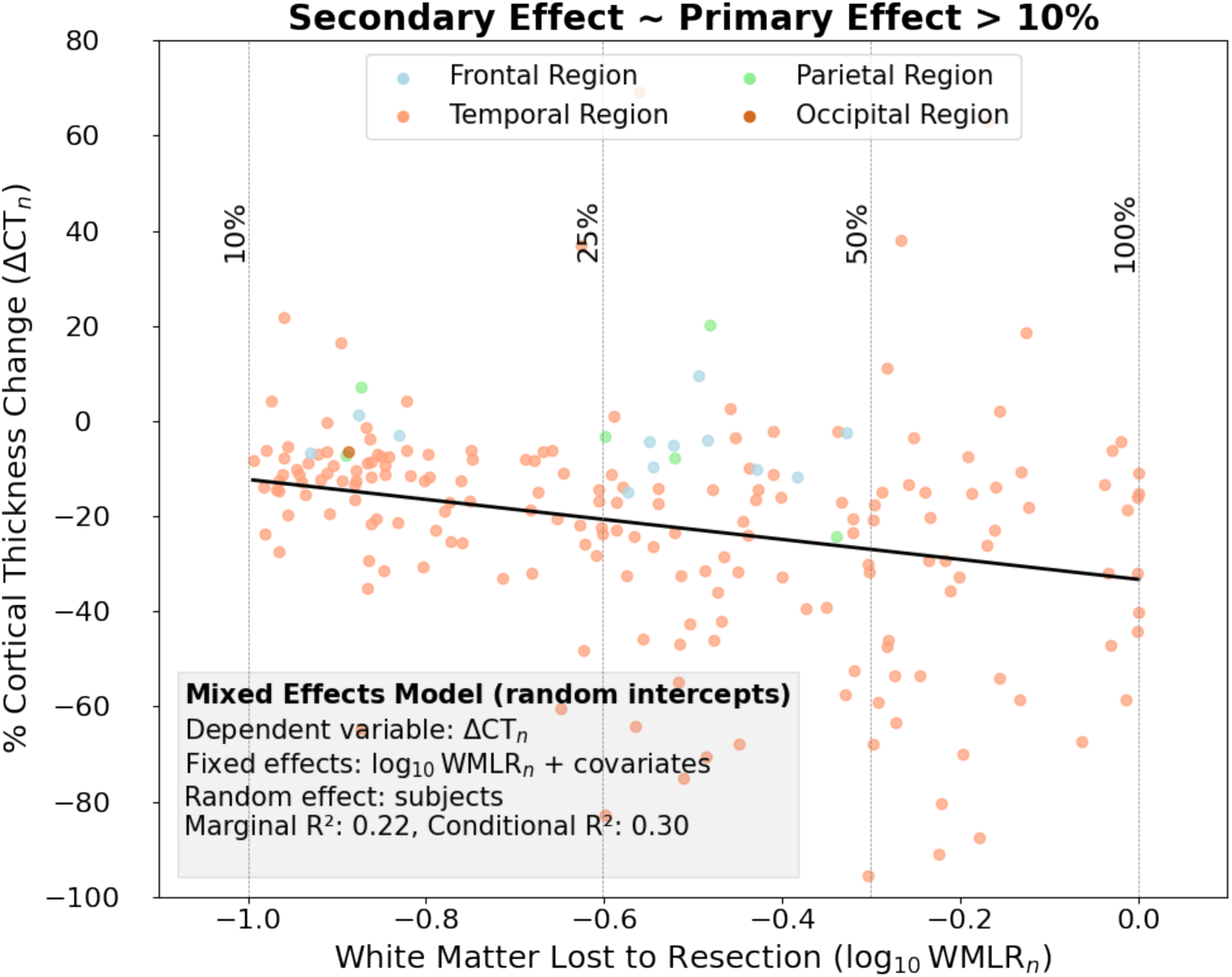
Relationship between the regional White Matter Lost to Resection and the regional change in Cortical Thickness for regions with white matter losses >10%. While the unthresholded model suggests a 3.4% decrease in cortical thickness for each 10-fold loss of white matter, the thresholded model suggests a 21% decrease in cortical thickness. This corresponds to 6.2 times accelerated cortical thinning for regions with white matter losses above this threshold.

### Towards comprehensive models of interventional consequence

The presented results challenge the notion that the prospective consequences of surgical white matter resection need only consider disruption of nearby eloquent bundles. We have shown not only that surgical white matter resection implicates a broad spectrum of interregional connections – impacting both regions close and remote to resection – but also that they are associated with manifest morphological changes that actively remodel brain anatomy. These impacts to both local circuitry and widespread brain networks may provide explanatory power where there are complicated spectra of currently unexplained postoperative deficits *(e.g., complex cognitive outcomes, mood disturbances, etc.)* despite preservation of such eloquent bundles.

Moving towards more comprehensive models of postoperative consequences may especially benefit the surgical treatment of epilepsy. In contrast to procedures for other neurological conditions, resection of white matter here also contributes to the intervention’s therapeutic effect, supporting seizure control through the disruption of propagation networks^29^. Building on this principle, a postoperative atrophy of non-resected propagation networks may help to further suppress seizure transmission^30^; a potential long-term recovery of atrophic networks^31^ could provide a structural explanation for late-onset seizure recurrence. For patients who continue to experience seizures post-resection, the altered brain network architecture could constitute a structural correlate for changed seizure semiology.

While the specific relationships between anatomical alterations and clinical outcomes will need to be investigated in future studies, comprehensive assessment of anatomical change holds significant potential to understand currently unexplained outcomes. Encouragingly, our parsimonious imaging approach allowed reliable modelling of degenerative effects through virtual resection of dMRI-based streamline tractograms. With accurate mapping of individual white matter and modern predictive modelling techniques, this raises the exciting prospect of integrating streamline tractography into surgical planning to predict postoperative atrophy prior to intervention.

### Network-wide structural plasticity may be more limited than previously thought

The brain has remarkable capabilities to functionally adapt to changes imposed by resections, e.g. shifting language functions to cortices adjacent to resection or even to the contralateral hemisphere^32,33^. In line with this notion of brain plasticity, observations of postoperatively increased white matter connectivity are often thought to be the structural basis for such functional adaptions; yet, there is a discordance between neuroanatomical understanding, where the adult human brain does not establish new long-range axonal connections^34^, and the results of network neuroscience studies, where gross post-surgical modulation of structural connectivity has been reported^35–37^. Given the considerable reconstruction variance of tractography even in scan-rescan experiments^38^, there is a concern that such extensive connectivity findings are *(at least partially)* the result of unaccounted methodological variance.

In fact, if we had estimated change in downstream connectivity using the conventional strategy of cross-sectionally reconstructing timepoints, reported results would have differed drastically, showing non-specific increases and decreases across both hemispheres *(see Extended Data Fig. 2A)*. In isolation, such data could reasonably have been interpreted as *“widespread white matter impairments”* accompanied by *“structural reorganisations”*, in agreement with similar reports in the literature^8,31,35–37,39,40^. However, the availability of an improved robust method that primarily yields connectivity decreases specific to the site of resection raises concerns about the validity of longitudinal differences estimated from cross-sectional reconstruction *(see Extended Data Fig. 2B)*. The concern is further strengthened when evaluating the relationship between our primary and tertiary effect: while the regional White Matter Lost to Resection *(primary effect)* was able to explain 58% of variance when change in Downstream Connectivity *(tertiary effect)* was estimated in a robust manner, it explained less than 1% of variance when it was estimated using cross-sectional methods *(see Extended Data Fig. 2C)*.

The failure to capture this biologically expected relationship, as well as the generally decreased anatomical specificity of cross-sectionally estimated connectivity changes, underscores the importance of mitigating reconstruction variance in longitudinal analyses. The fact that gross connectivity increases were not replicated using robust longitudinal methods suggests them to be methodological artefacts rather than true biological phenomena – meaning that the human brain’s capacity for postoperative white matter plasticity may be more limited than previously suggested.

Although we did not find evidence to support network-wide reorganisations, our robust cortical thickness analysis demonstrated hypertrophic cortices in both parietal lobes and lateral occipital cortices after tsSAHE. The observation of limited cortical thickness increases following restricted resections is consistent with recent work employing robust methods, demonstrating cortical thickness increases only within the supramarginal gyrus after *transcortical* SAHE, yet not after ATL^2^. However, in the absence of a clear *a priori* hypothesis we are cautious ascribing these findings to local structural adaptations, as subtle cortical hypertrophy may also be epilepsy-related *(e.g. to changed anti-seizure medication)*. Further studies should look into the microstructural changes within hypertrophic cortices to better characterise the biological nature of underlying processes as well as their clinical relevance.

**Extended Data Figure 2.**
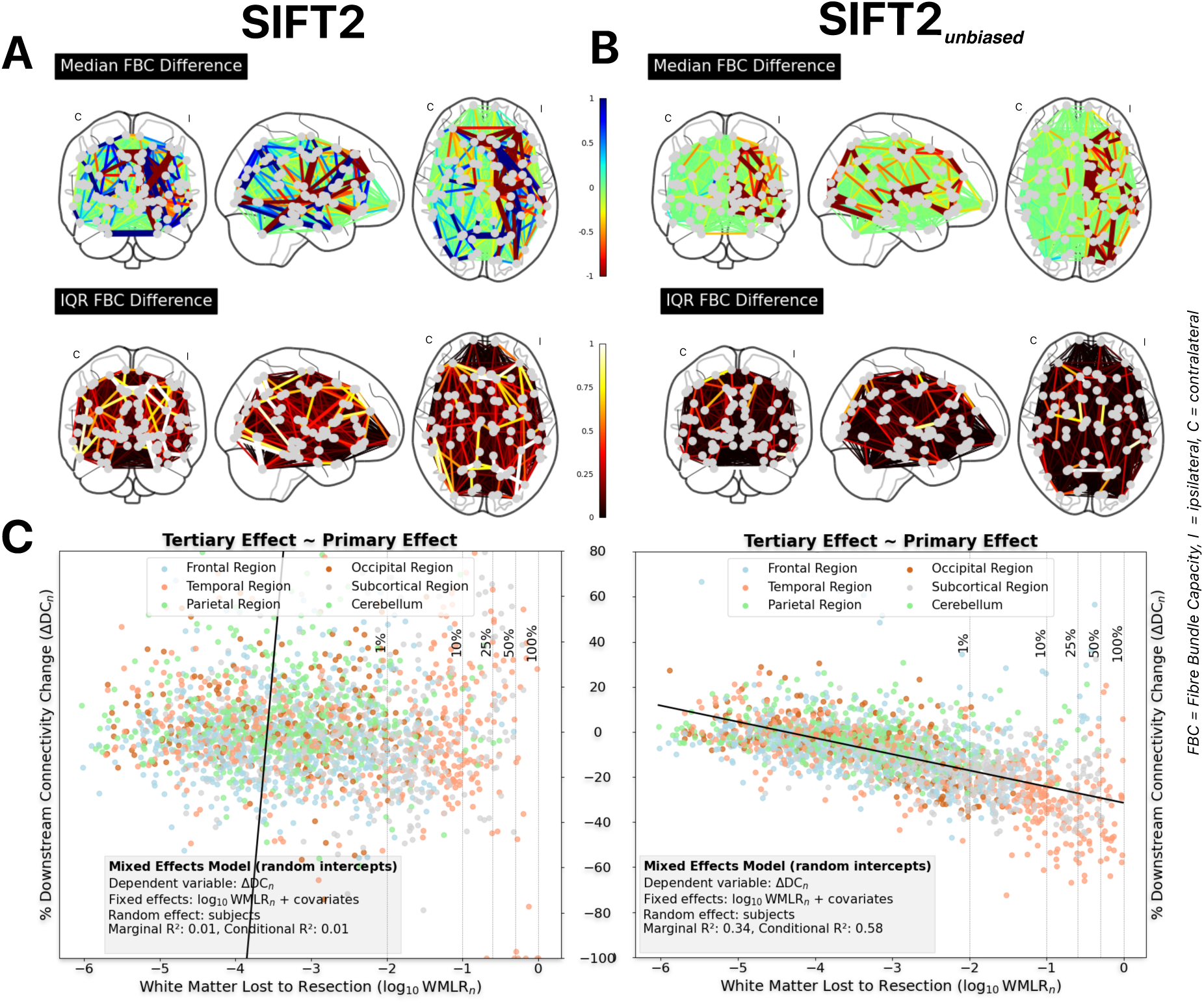
SIFT2 vs. SIFT2_unbiased_ for longitudinal connectivity analysis. A) Across our cohort, cross-sectional reconstruction led to non-specific longitudinal increases and decreases in Fibre Bundle Capacity (FBC), with increased edge-wise standard deviations. B) Unbiased reconstruction on the other hand delineated FBC decreases mainly within the hemisphere ipsilateral to resection, as would be biologically expected. C) Evaluating the relationships between surgically resected white matter and independently estimated downstream connectivity change explained less than 1% of data variance. If downstream connectivity change was estimated in a robust manner, the model explained 58% of variance. ATL = anterior temporal lobectomy, C = contralateral, FBC = Fibre Bundle Capacity, tsSAHE = transsylvian selective amygdalohippocampectomy, I = ipsilateral, IQR = interquartile range.

## Limitations

Our postoperative data was restricted to the period approximately six months following resection; while this relatively short timeframe is ideal to isolate the effects of transneuronal degeneration by minimising the influence of external factors *(e.g. changes in medication or postoperative seizure-recurrence),* future studies should clarify the long-term anatomical effects of resective surgery. The dMRI protocols used for this study adhere to clinical standards, however, advanced multi-shell sequences with high spatial and angular contrast would enhance accuracy of white matter reconstructions. This would allow improved characterisation of the biologically expected relationships between surgically resected white matter and downstream anatomical changes. Nonetheless, it is highly encouraging that our longitudinal analysis approach derived biologically plausible connectivity estimates from standard clinical protocols.

The effects of transneuronal degeneration were here demonstrated exclusively based on neuroanatomical and not clinical consequences. The latter would provide corroborating motivation for the clinical adoption of advanced tractography to anticipate interventional consequences but was neither within the scope of this paper nor possible with our exemplar cohort.

Lastly, we modelled postoperative change as a consequence of resected white matter, but other mechanisms, e.g. intraoperative tissue retraction or microvascular damage, may also induce anatomical alterations. While such alternative mechanisms could not be assessed using methods employed for this study, they may help to account for variance currently unexplained by our model.

## Conclusion

This imaging study provides compelling support that degenerative effects of neurosurgery can propagate via white matter, demonstrating a sequential network effect concordant with the mechanism of *transneuronal degeneration*. Our longitudinal imaging analysis framework is tailored for the exploration of this effect and can be readily utilised to explore it across other neurological diseases. Evidence of this mechanism across the brain’s ubiquitous neural networks raises the prospect of utilising advanced dMRI-based tractography to predict interventional consequences arising from the disruption of axonal pathways.

## Methods

### Inclusion criteria and clinical workflow

This study included temporal lobe epilepsy patients who underwent ATL or tsSAHE between 2012 and 2019 at the Medical University of Vienna, with available pre- and postoperative diffusion-weighted and T1-weighted MRI sequences. A detailed description of the clinical workflow as well as technical aspects the surgical procedures is published elsewhere^13^ and is reproduced in the *Supplementary Materials*.

In brief, seizure onset and laterality were confirmed by preoperative video-EEG-monitoring by experienced epileptologists, followed by a comprehensive preoperative workup that included a standardised epilepsy imaging protocol at 3T *(Philips Achieva).* Individual findings were discussed in an interdisciplinary epilepsy board to determine eligibility for surgery, with patients that had clear signs of a mesial temporal lobe seizure onset undergoing tsSAHE, patients with unclear mesial temporal signs, or additional extramesial structural changes undergoing ATL. All surgeries were performed by the same experienced epilepsy surgeon. ATL was performed as standard ATL *(en-bloc resection of the anterior temporal lobe; 3-4cm in the dominant, 4-5cm in the non-dominant hemisphere)*^22^, with one patient receiving a left-sided hippocampal-sparing resection. For all SAHE procedures, the transylvian-transamygdala approach as originally described by Yasargil et. al^19–21^ was followed.

Altogether, a series of 79 consecutively operated patients *(ATL:SAHE=43:36)* was identified, of which 59 patients *(ATL:SAHE=31:28)* fulfilled the inclusion criteria of this study. Eleven ATL patients were operated within the left and 20 ATL patients within the right hemisphere; for patients that underwent tsSAHE, 16 were operated within the left and 12 were operated within the right hemisphere. No significant difference in baseline characteristics was found between treatment groups, except that more male patients underwent ATL *(see Table 1 and Supplementary Materials)*.

**Table 1.**
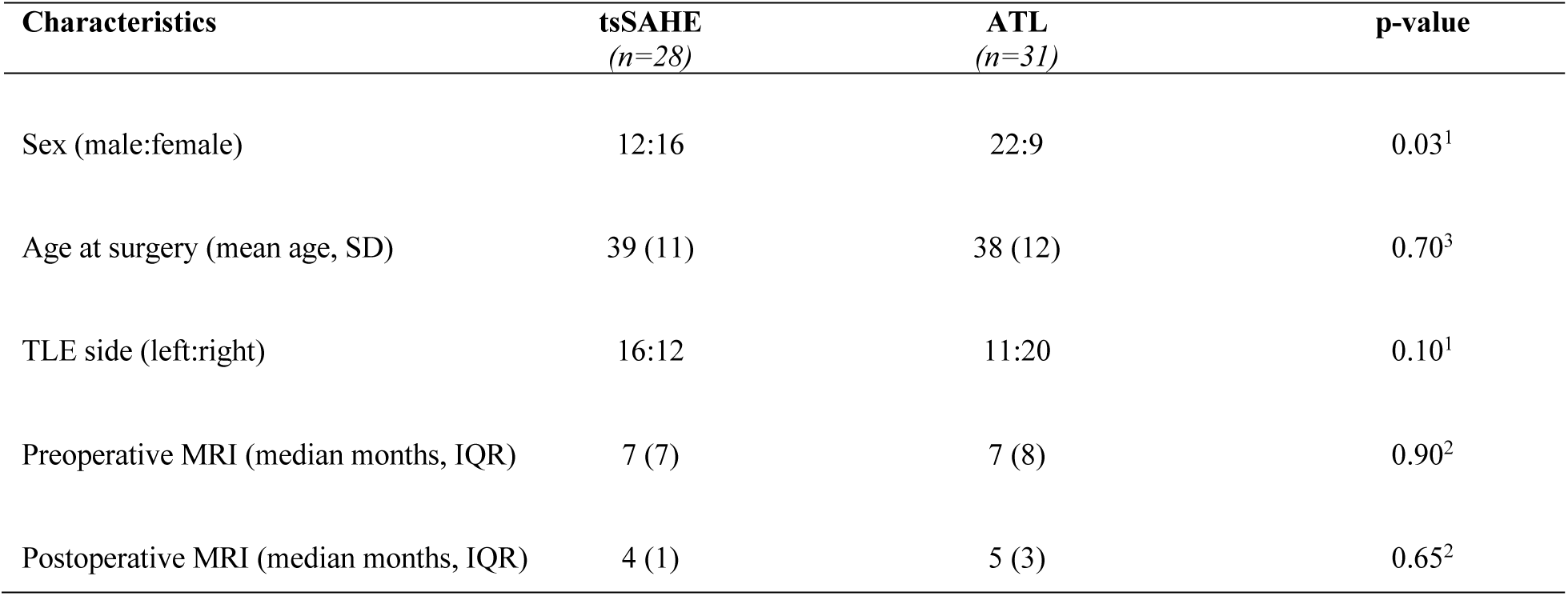
Baseline Characteristics. We found no differences between treatment groups, except that male patients underwent ATL more frequently. A comprehensive overview of the cohort’s characteristics can be found in the Supplementary Materials. ATL = anterior temporal lobectomy; IQR = interquartile range; SD = standard deviation; TLE = temporal lobe epilepsy, tsSAHE = transsylvian selective amygdalohippocampectomy; Statistical Tests: 1= Qui-squared test, 2 = Mann-Whitney U test, 3 = Student’s t-test.

### Imaging sequences

Patients underwent a standardised epilepsy protocol before and after surgery, acquired on a 3T MRI scanner *(Achieva, Philips Healthcare Systems)*.

Pre- and postoperative diffusion-weighted images were acquired with the following parameters: voxel size = 2mm^3^ isotropic, number of slices = 60, matrix size = 128×128, field of view = 256×256mm, repetition time = 6454ms, echo time = 70ms, in-plane parallel acceleration = 2x, partial Fourier = 0.86, number of non-collinear gradients = 32, *b*-value = 800 s/mm^2^, number reference images without diffusion weighting *(b=0)* = 1.

Pre- and postoperative T1w 3D FLASH images were acquired with the following parameters: voxel size = 1mm^3^ isotropic, number of slices = 190, matrix = 240×240, field of view = 240×240mm, repetition time = 8.1ms, echo time = 3.7ms.

For 13 patients *(ATL:SAHE=3:4)* one of the acquired T1w images differed from the standardised protocol, but were included into longitudinal analysis after assessing the risk of asymmetry bias^41^ (see *Supplementary Materials*).

### Expected imaging effects following surgical tissue resection

Following surgical tissue resection, we expected three structural effects to sequentially manifest for non-resected brain regions, namely the (1) loss of surgically disrupted axons, (2) the deterioration of disconnected neurons as well as (3) the degeneration of axonal projections thereof. Using advanced neuroimaging reconstructions, we evaluated the presence of these effects as follows:

1. **Primary Effect: Regional White Matter Lost to Resection.** The extent of white matter involved in resection for a given brain region, determined by the fraction of cross-sectional weights of tractography streamlines that intersected the resection cavity. Cross-sectional weights were estimated with SIFT2^24^.
2. **Secondary Effect: Regional Cortical Thickness Change.** The fractional change in cortical thickness for a given brain region, estimated within a longitudinal Freesurfer^27,28,41,42^ framework adapted for the use of surgical image data.
3. **Tertiary Effect: Regional Downstream Connectivity Change.** The change in connection density within non-resected white matter projecting to/from a given brain region, determined by the fractional change in cross-sectional weights of tractography streamlines that did not intersect the resection cavity. Cross-sectional weights were longitudinally estimated within the robust SIFT2_unbiased_ framework specifically designed for this study.

For regions that underwent partial resection, metrics were only quantified for surgically preserved parts of the regions *(see Fig 3B).* Through these regional effects, we constructed an imaging model of transneuronal degeneration, employing two linear mixed effects models that evaluated the explanatory power of surgically resected white matter *(primary effect)* for downstream anatomical changes *(secondary and tertiary effects)*.

### Image processing

Image processing was performed using Neurodesk^43^, a containerised environment for reproducible neuroimaging. Pre-processing of diffusion data was performed using MRtrix3^44^ *(v3.0.3)*, FSL *(v6.0.5.1)*, SynB0-DISCO *(v3.0)*, and ANTs^45^ *(v2.3.5)*. Further diffusion analysis was performed using MRtrix3 *(v.3.0.3)*^44^ or augmentations thereof. Detailed descriptions of individual processing steps are provided in the *Supplementary Materials*.

In short, diffusion data was denoised^46^ and corrected for Gibbs ringing^47^, susceptibility distortions and eddy currents^48,49^ as well as B1 field inhomogeneities^50^. Note that for susceptibility distortion correction we used a synthetic *b=*0 image, generated using synb0-DISCO^51^. Macroscopic tissue response functions were estimated from preoperative diffusion data using a data-driven approach^52^ and averaged across participants. We performed Single-Shell 3-Tissue *(SS3T)* constrained spherical deconvolution^53^ to estimate orientation distribution functions *(ODFs)* that were then intensity normalised^54^. Structural data were processed using Freesurfer^26^ *(v7.4.1)*, including N3 bias field correction^55^, intensity normalisation^56^ and were then rigidly registered to preprocessed diffusion images using ANTs^45^ *(v2.3.5)*.

### Template generation and resection cavity segmentation

To aggregate preoperative and postoperative image data, we generated subject-specific unbiased^41^ templates across sessions. First, structural images were rigidly aligned using robust template registration^42^, followed by symmetric non-linear refinement^45^. The resection cavity was segmented from transformed structural images using a fully automated segmentation approach *(see Supplementary Materials).* Finally, unbiased diffusion templates were generated by averaging transformed, reoriented and modulated^58^ fiber ODFs.

Note that for areas of tissue resection, image information exclusively was contributed by preoperative sessions, allowing reconstruction of anatomically intact average timepoints *(see Supplementary Fig. 2)*.

### An adapted longitudinal FreeSurfer framework for use of surgical image data

FreeSurfer’s longitudinal processing stream is an established framework for robust longitudinal estimation of cortical thickness^26,27^. Surgical data however violates its assumption of fixed gross anatomy over time. Therefore, we adapted the existing Freesurfer longitudinal pipeline to suit data involving gross resection as follows: after initialising the longitudinal processing stream, we injected the previously generated *(non-linearly refined and anatomically corrected)* subject template, allowing reconstruction of an unbiased representative yet anatomically intact subject template^28^. This information was then transferred back to individual timepoints to initialise session-specific optimisation as per longitudinal Freesurfer pipeline design. Cortical vertices were individually tested for intersection with the resection volume in subject template space. Cortical parcels were defined based on the default Desikan-Killiany^23^ parcellation; parcels with partial intersection with the subject resection were bisected into resected and preserved portions, with longitudinal quantitative measures calculated exclusively from the latter. One subject was excluded from cortical thickness analysis due to the presence of a postoperative subdural hematoma that prevented reliable parameter estimation.

### SIFT2_unbiased_: robust estimation of longitudinal brain connectivity

The adult human brain has very limited capacity to form new axonal connections^34^, allowing robust estimation of longitudinal connectivity to follow the same strategy as longitudinal FreeSurfer: reconstruction of gross anatomy from an unbiased, representative template^41^, which can be then refined to fit individual timepoints.

For this study, we constructed base streamline tractograms from unbiased fiber ODF templates *(10 million streamlines, dynamic seeding*^24^*, anatomically constrained tractography*^59^) and estimated corresponding connection densities using SIFT2^24^. These weighted tractograms were then transferred to individual timepoints for further refinement using SIFT2_unbiased_, returning connection densities within previously constructed tractograms that fit individual session fiber densities. Note that surgical data also in this case violates the underlying assumption of fixed gross anatomy; therefore, before initialising postoperative refinement, we set all density-weights of streamlines intersecting the resection mask to zero, effectively excluding resected connections from optimisation.

Finally, streamlines were assigned to previously obtained parcels and connection densities summarised to *region-wise* connectomes, computing the total pre- and postoperative white matter density for each surgically preserved region. Connections were multiplied by the corresponding proportionality coefficient *μ* to appropriately scale densities between sessions^25^. The requisite code to run this pipeline is distributed via the MRtrix3 software package (https://github.com/MRtrix3/mrtrix3/tree/tcksift2_weight_options). Further details about the SIFT2_unbiased_ framework can be found in the *Supplementary Materials*.

### Comparison of robust versus non-robust longitudinal estimation of connectivity

To demonstrate the benefits of robust longitudinal connectivity analysis *(SIFT2_unbiased_)* versus non-robust longitudinal connectivity analysis *(cross-sectional SIFT2),* we additionally generated density weighted tractograms using the same software outside our longitudinal framework.

For each participant, we cross-sectionally constructed pre- and postoperative streamline tractograms from session fiber ODF data using the same tracking parameters as were used for unbiased reconstruction *(10 million streamlines, dynamic seeding*^24^*, anatomically constrained tractography*^59^). Streamline densities were optimised to match their respective session fiber ODF data using SIFT2^24^.

Streamlines of all generated tractograms were assigned to non-resected parts of brain regions and streamline densities summarised to *edge-wise* connectomes. Each connectome edge was multiplied by the corresponding proportionality coefficient *μ* to compute the pre- and postoperative Fibre Bundle Capacity between surgically preserved regions^25^.

An overview of the differences of robustly versus non-robustly estimated connectivity results is demonstrated in *Extended Data Fig. 3* and in the *Supplementary Materials*.

### Quantification of Primary Effect: Regional White Matter Lost to Resection

The regional White Matter Lost to Resection *(primary effect)* was quantified for non-resected parts of brain regions as:

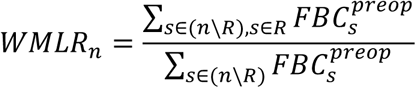

where *WMLR_n_* is the fraction of white matter lost to resection for region *n*, 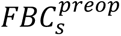 is the Fibre Bundle Capacity^25^ ascribed to streamline *s* in the preoperative data, *s∈(n\R)* indicates that streamline *s* is ascribed to the portion of region *n* that does not intersect the resection region *R*, and *s∈R* indicates that streamline *s* intersects resection cavity *R*.

### Quantification of Secondary Effect: Regional Cortical Thickness Change

The regional change in Cortical Thickness *(secondary effect)* was quantified longitudinally for non-resected parts of brain regions as:

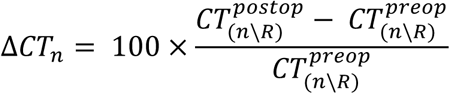

where Δ*CT_n_* is the fractional change in cortical thickness for region *n*, (*n* ∖ *R*) indicates the portion of region *n* that does not intersect resection region 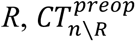 and 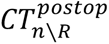 are the mean cortical thickness in this portion of region *n* for preoperative and postoperative data respectively.

### Quantification of Tertiary Effect: Regional Downstream Connectivity Change

The regional change in Downstream Connectivity *(tertiary effect)* was quantified longitudinally for non-resected parts of brain regions as:

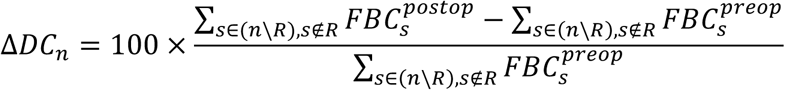

where Δ*DC_n_* is the change in downstream white matter connectivity for region *n*, *s* ∈ (*n*\*R*) indicates that streamline *s* is ascribed to the portion of region *n* that does not intersect resection *R*, *s* ∉ R indicates that streamline *s* does not intersect resection region *R*, and 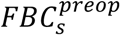 and 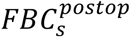 indicate the Fibre Bundle Capacity^25^ ascribed to streamline *s* in the preoperative and postoperative session data respectively.

### Reporting, statistical testing and false-positive control

White Matter Lost to Resection *(primary effect)* was reported if observed losses were ≥ 1%; losses below this threshold were considered clinically negligible. Secondary and tertiary effects were tested for statistical longitudinal differences *(scipy*^60^, *v1.11.4)* using a paired t-test and considered statistically significant with a two-sided significance level of alpha <0.05 after correction for multiple comparisons *(False-Discovery-Rate)*.

### Imaging model of transneuronal degeneration

We used mixed effects models *(statsmodels*^61^*, v.0.14.2)* with random intercepts to investigate the explanatory power of surgically resected white matter for neuroanatomical changes downstream of resection, namely changes in cortical thickness and downstream connectivity. Relationships were explored using brain regions that had streamline intersections with the resection cavity, which were pooled across all subjects.

In particular, we used the following model to predict longitudinal change in Cortical Thickness:

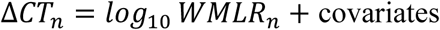

where Δ*CT_n_* is the longitudinal change in Cortical Thickness for a region *n (secondary effect)* and *log*_10_ *WMLR_n_* is the log_10_-transformed White Matter Lost to Resection for a region *n (primary effect)*.

To predict longitudinal change in Downstream Connectivity, we used the following model

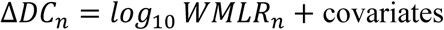

where Δ*DC_n_* is the longitudinal change in Downstream Connectivity for a region *n (tertiary effect)* and *log*_10_ *WMLR_n_* is the log_10_-transformed White Matter Lost to Resection for a region *n (primary effect)*.

For both models, clinical covariates were included as fixed effects, namely treatment group, side of surgery, sex, disease duration, and age at surgery. Subjects were included as random effects to account for the hierarchical data structure. Note that disease duration and age at surgery were missing for one patient that underwent ATL *(lost to follow up);* to include the subject into the regression model, missing covariates were imputed for this subject based on the cohort mean.

### Ethic Approval

This study was approved by the Ethics Committee of the Medical University of Vienna (Study Number 2135/2021). It was conducted in line with the Declaration of Helsinki.

## Abbreviations

ATL: Anterior Temporal Lobectomy
ΔCT_n_: Cortical Thickness Change for brain region n (quantitative imaging parameter)
ΔDC_n_: Downstream Connectivity Change for brain region n (quantitative imaging parameter)
(d)MRI: (diffusion) Magnetic Resonance Imaging
FBC: Fibre Bundle Capacity
ODF: Orientation Distribution Function
IQR: Interquartile Range
SIFT: Spherical Deconvolution Informed Filtering of Tractograms
(ts)SAHE: (transsylvian) Selective Amygdalohippocampectomy
WMLR_n_: White Matter Lost to Resection for brain region n (quantitative imaging parameter)

## Acknowledgments

P.P is supported by the Melbourne International Research Scholarship from the University of Melbourne. R.M. is the recipient of an Australian Research Council Discovery Early Career Researcher Award *(project number: DE240101035).* R.E.S. is supported by fellowship funding from the National Imaging Facility *(NIF),* an Australian Government National Collaborative Research Infrastructure Strategy *(NCRIS)* capability. The work was further supported by the Developmental and Interventional Neuroimaging Lab *(DINLAB)* at the Division of Neuroradiology and Musculoskeletal Radiology of the Department of Biomedical Imaging and Image-guided Therapy at the Medical University of Vienna. We also thank the Statistical Consulting Centre *(School of Mathematics and Statistics)* at the University of Melbourne for advising on the statistical design of this study.

## References

1. Dewan, M. C. et al. Global neurosurgery: the current capacity and deficit in the provision of essential neurosurgical care. Executive Summary of the Global Neurosurgery Initiative at the Program in Global Surgery and Social Change. Journal of Neurosurgery 130, 1055–1064 (2019).

2. Arnold, T. C. et al. Remote effects of temporal lobe epilepsy surgery: Long-term morphological changes after surgical resection. Epilepsia Open 8, 559–570 (2023).

3. Leiberg, K. et al. Effects of anterior temporal lobe resection on cortical morphology. Cortex 166, 233–242 (2023).

4. Lam, J. et al. Gray Matter Atrophy: The Impacts of Resective Surgery and Vagus Nerve Stimulation in Drug-Resistant Epilepsy. World Neurosurgery 149, e535–e545 (2021).

5. Elliott, C. A., Gross, D. W., Wheatley, B. M., Beaulieu, C. & Sankar, T. Progressive contralateral hippocampal atrophy following surgery for medically refractory temporal lobe epilepsy. Epilepsy Research 125, 62–71 (2016).

6. Schoene-Bake, J.-C. et al. Widespread affections of large fiber tracts in postoperative temporal lobe epilepsy. NeuroImage 46, 569–576 (2009).

7. McDonald, C. R. et al. Changes in fiber tract integrity and visual fields after anterior temporal lobectomy. Neurology 75, 1631–1638 (2010).

8. Faber, J. et al. Progressive fiber tract affections after temporal lobe surgery. Epilepsia 54, e53–e57 (2013).

9. Waller, A. V. & Owen, R. Experiments on the section of the glossopharyngeal and hypoglossal nerves of the frog, and observations of the alterations produced thereby in the structure of their primitive fibres. Abstracts of the Papers Communicated to the Royal Society of London 5, 924–925 (1997).

10. Cowan, W. M. Anterograde and Retrograde Transneuronal Degeneration in the Central and Peripheral Nervous System. in Contemporary Research Methods in Neuroanatomy (eds. Nauta, W. J. H. & Ebbesson, S. O. E.) 217–251 (Springer Berlin Heidelberg, Berlin, Heidelberg, 1970). doi:10.1007/978-3-642-85986-1_11.

11. Fornito, A., Zalesky, A. & Breakspear, M. The connectomics of brain disorders. Nature Reviews Neuroscience 16, 159–172 (2015).

12. Binding, L. P. et al. Contribution of White Matter Fiber Bundle Damage to Language Change After Surgery for Temporal Lobe Epilepsy. Neurology 100, e1621–e1633 (2023).

13. Pruckner, P. et al. Visual outcomes after anterior temporal lobectomy and transsylvian selective amygdalohippocampectomy: A quantitative comparison of clinical and diffusion data. Epilepsia 64, 705–717 (2023).

14. Schilling, K. G. et al. Short superficial white matter and aging: A longitudinal multi-site study of 1293 subjects and 2711 sessions. Aging Brain 3, 100067 (2023).

15. Beatty, R., Sadun, A., Smith, L., Vonsattel, J. & Richardson, E. Direct demonstration of transsynaptic degeneration in the human visual system: a comparison of retrograde and anterograde changes. *Journal of Neurology*, Neurosurgery & Psychiatry 45, 143–146 (1982).

16. Cowey, A., Alexander, I. & Stoerig, P. Transneuronal retrograde degeneration of retinal ganglion cells and optic tract in hemianopic monkeys and humans. Brain 134, 2149–2157 (2011).

17. Jindahra, P., Petrie, A. & Plant, G. T. The time course of retrograde trans-synaptic degeneration following occipital lobe damage in humans. Brain 135, 534–541 (2012).

18. Gupta, N. et al. Atrophy of the lateral geniculate nucleus in human glaucoma detected by magnetic resonance imaging. Br J Ophthalmol 93, 56 (2009).

19. Dorfer, C. et al. Mesial temporal lobe epilepsy: long-term seizure outcome of patients primarily treated with transsylvian selective amygdalohippocampectomy. Journal of neurosurgery 129, 174–181 (2017).

20. Yaşargil, M. G., Türe, U. & Yaşargil, D. C. Impact of temporal lobe surgery. Journal of neurosurgery 101, 725–738 (2004).

21. Yasargil, M., Teddy, P. & Roth, P. Selective amygdalo-hippocampectomy operative anatomy and surgical technique. Advances and Technical Standards in Neurosurgery: Volume 2 93–123 (1985).

22. Zentner, J. Surgical Treatment of Epilepsies. (Springer, 2002).

23. Desikan, R. S. et al. An automated labeling system for subdividing the human cerebral cortex on MRI scans into gyral based regions of interest. NeuroImage 31, 968–980 (2006).

24. Smith, R. E., Tournier, J.-D., Calamante, F. & Connelly, A. SIFT2: Enabling dense quantitative assessment of brain white matter connectivity using streamlines tractography. NeuroImage 119, 338–351 (2015).

25. Smith, R. E., Raffelt, D., Tournier, J.-D. & Connelly, A. Quantitative streamlines tractography: methods and inter-subject normalisation. Aperture Neuro 1–25 (2022).

26. Fischl, B. FreeSurfer. NeuroImage 62, 774–781 (2012).

27. Reuter, M., Schmansky, N. J., Rosas, H. D. & Fischl, B. Within-subject template estimation for unbiased longitudinal image analysis. NeuroImage 61, 1402–1418 (2012).

28. Hoffmann, M., Salat, D., Reuter, M. & Fischl, B. Longitudinal FreeSurfer with non-linear subject-specific template improves sensitivity to cortical thinning. Athinoula A. Martinos Center for Biomedical Imaging, Charlestown, MA, United States; Department of Radiology, Harvard Medical School, Boston, MA, United States; German Center for Neurodegenerative Diseases, Bonn, Germany; Computer Science and Artificial Intelligence Laboratory, Massachusetts Institute of Technology, Cambridge, MA, United States.

29. Neal, E. G., Maciver, S., Schoenberg, M. R. & Vale, F. L. Surgical disconnection of epilepsy network correlates with improved outcomes. Seizure 76, 56–63 (2020).

30. Jirsa, V. K. et al. The Virtual Epileptic Patient: Individualized whole-brain models of epilepsy spread. NeuroImage 145, 377–388 (2017).

31. Li, W. et al. Different patterns of white matter changes after successful surgery of mesial temporal lobe epilepsy. NeuroImage: Clinical 21, 101631 (2019).

32. Nieberlein, L., Rampp, S., Gussew, A., Prell, J. & Hartwigsen, G. Reorganization and Plasticity of the Language Network in Patients with Cerebral Gliomas. NeuroImage: Clinical 37, 103326 (2023).

33. Duffau, H. Lessons from brain mapping in surgery for low-grade glioma: insights into associations between tumour and brain plasticity. The Lancet Neurology 4, 476–486 (2005).

34. Cafferty, W. B. J., McGee, A. W. & Strittmatter, S. M. Axonal growth therapeutics: regeneration or sprouting or plasticity? Trends in Neurosciences 31, 215–220 (2008).

35. Jeong, J.-W., Asano, E., Juhász, C., Behen, M. E. & Chugani, H. T. Postoperative axonal changes in the contralateral hemisphere in children with medically refractory epilepsy: A longitudinal diffusion tensor imaging connectome analysis. Human Brain Mapping 37, 3946–3956 (2016).

36. Ji, G.-J. et al. Connectome Reorganization Associated With Surgical Outcome in Temporal Lobe Epilepsy. Medicine 94, (2015).

37. Larivière, S. et al. Connectome reorganization associated with temporal lobe pathology and its surgical resection. Brain 147, 2483–2495 (2024).

38. Smith, R. E., Tournier, J.-D., Calamante, F. & Connelly, A. The effects of SIFT on the reproducibility and biological accuracy of the structural connectome. NeuroImage 104, 253–265 (2015).

39. da Silva, N. M. et al. Network reorganisation following anterior temporal lobe resection and relation with post-surgery seizure relapse: A longitudinal study. NeuroImage: Clinical 27, 102320 (2020).

40. Yogarajah, M. et al. The structural plasticity of white matter networks following anterior temporal lobe resection. Brain 133, 2348–2364 (2010).

41. Reuter, M. & Fischl, B. Avoiding asymmetry-induced bias in longitudinal image processing. NeuroImage 57, 19–21 (2011).

42. Reuter, M., Rosas, H. D. & Fischl, B. Highly accurate inverse consistent registration: A robust approach. NeuroImage 53, 1181–1196 (2010).

43. Renton, A. I. et al. Neurodesk: an accessible, flexible and portable data analysis environment for reproducible neuroimaging. Nature Methods 21, 804–808 (2024).

44. Tournier, J.-D. et al. MRtrix3: A fast, flexible and open software framework for medical image processing and visualisation. NeuroImage 202, 116137 (2019).

45. Avants, B. B. et al. A reproducible evaluation of ANTs similarity metric performance in brain image registration. NeuroImage 54, 2033–2044 (2011).

46. Veraart, J. et al. Denoising of diffusion MRI using random matrix theory. NeuroImage 142, 394–406 (2016).

47. Kellner, E., Dhital, B., Kiselev, V. G. & Reisert, M. Gibbs-ringing artifact removal based on local subvoxel-shifts. Magnetic Resonance in Medicine 76, 1574–1581 (2016).

48. Smith, S. M. et al. Advances in functional and structural MR image analysis and implementation as FSL. NeuroImage 23, S208–S219 (2004).

49. Andersson, J. L. R. & Sotiropoulos, S. N. An integrated approach to correction for off-resonance effects and subject movement in diffusion MR imaging. NeuroImage 125, 1063–1078 (2016).

50. N. J. Tustison et al. N4ITK: Improved N3 Bias Correction. IEEE Transactions on Medical Imaging 29, 1310–1320 (2010).

51. Schilling, K. G. et al. Synthesized b0 for diffusion distortion correction (Synb0-DisCo). Magnetic Resonance Imaging 64, 62–70 (2019).

52. Dhollander, T., Raffelt, D. & Connelly, A. Unsupervised 3-Tissue Response Function Estimation from Single-Shell or Multi-Shell Diffusion MR Data without a Co-Registered T1 Image. (2016).

53. Dhollander, T., Mito, R., Raffelt, D. & Connelly, A. Improved White Matter Response Function Estimation for 3-Tissue Constrained Spherical Deconvolution. (2019).

54. Raffelt, D., et al. Bias Field Correction and Intensity Normalisation for Quantitative Analysis of Apparent Fibre Density. (2017).

55. J. G. Sled, A. P. Zijdenbos, & A. C. Evans. A nonparametric method for automatic correction of intensity nonuniformity in MRI data. IEEE Transactions on Medical Imaging 17, 87–97 (1998).

56. Dale, A. M., Fischl, B. & Sereno, M. I. Cortical Surface-Based Analysis: I. Segmentation and Surface Reconstruction. NeuroImage 9, 179–194 (1999).

57. Raffelt, D., Tournier, J.-D., Crozier, S., Connelly, A. & Salvado, O. Reorientation of fiber orientation distributions using apodized point spread functions. Magnetic Resonance in Medicine 67, 844–855 (2012).

58. Raffelt, D. et al. Apparent Fibre Density: A novel measure for the analysis of diffusion-weighted magnetic resonance images. NeuroImage 59, 3976–3994 (2012).

59. Smith, R. E., Tournier, J.-D., Calamante, F. & Connelly, A. Anatomically-constrained tractography: Improved diffusion MRI streamlines tractography through effective use of anatomical information. NeuroImage 62, 1924–1938 (2012).

60. Virtanen, P. et al. SciPy 1.0: fundamental algorithms for scientific computing in Python. Nature Methods 17, 261–272 (2020).

61. Seabold, S. & Perktold, J. Statsmodels: Econometric and Statistical Modeling with Python. in *Proceedings of the 9th Python in Science Conference* (eds. Walt, S. van der & Millman, J.) 92–96 (2010). doi:10.25080/Majora-92bf1922-011.

