## Supplementary Materials for "Beyond the Resection: Surgical White Matter Disruption Structurally Alters Non-Resected Brain Anatomy"

#### SUPPLEMENTARY METHODS - MRI Acquisition Sequences

Analyses of this cohort have been previously published[1] and used the following MRI sequences:

***Diffusion weighted imaging***

Pre- and postoperative diffusion-weighted imaging (DWI) scans were acquired on the same 3T magnetic resonance imaging *(MRI)* scanner *(Achieva, Philips Healthcare Systems)* and were used for white matter reconstruction. Scans were obtained using the following imaging parameters: Voxel size: 2mm$^3$ isotropic, matrix size: 128x128, FOV: 256x256 mm, repetition time: 6454ms, echo time: 70ms, 60 slices, 2x parallel acceleration, partial Fourier: 0.86, number of non-collinear gradients: 32, b-value: 800 s/mm$^2$; additional acquisition of a reference image with no diffusion gradient applied ($b$=0).

***T1 weighted imaging***

Pre- and postoperative T1w images were acquired on the same 3T magnetic resonance imaging *(MRI)* scanner *(Achieva, Philips Healthcare Systems)* using a 3D FLASH sequence. Acquired T1w images were used for tissue segmentation, co-registration, template generation, and anatomical reconstruction. Scans were obtained using the following imaging parameters: Voxel size: 1mm$^3$ isotropic, matrix: 240x240, field of view *(FOV)*: 240x240 mm, repetition time: 8.1ms, echo time: 3.7ms, 190 slices.

##### *Impact of variation in T1 weighted acquisition protocols*

For a subset of patients, the acquired T1-weighted images did not conform to the standardised epilepsy protocol described above *(n=20)*. Subjects acquired with a protocol not present in at least 3 subjects were excluded from longitudinal cortical thickness assessment *(excluding 7 subjects)*. Of those remaining, most of these protocol variations occurred for postoperative scans only, introducing a risk of an asymmetry bias in quantification of longitudinal cortical thickness changes. To assess the risk of such bias, we systematically investigated the similarity of pre- and postoperative timepoints across the contralateral hemisphere, where biologically limited change is expected: no systematic outliers were detected following visual inspection *(see Supplementary Figure 1)*. Hence, these subjects were included in the longitudinal analysis of cortical thickness.

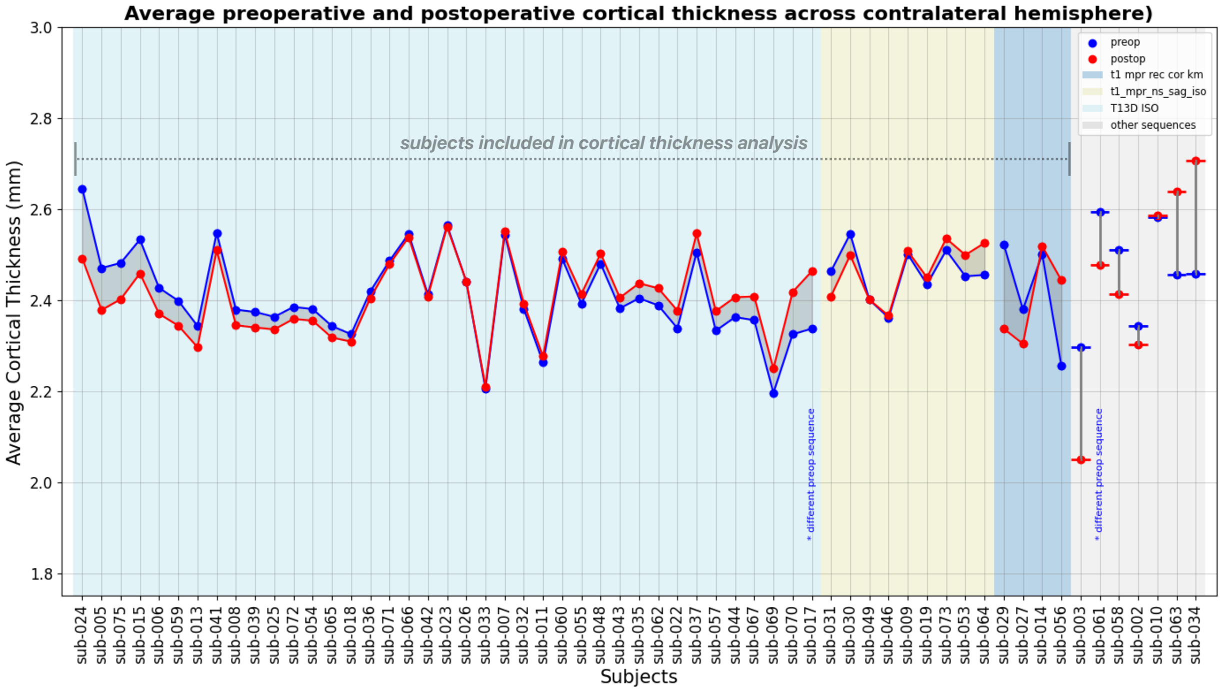


***Supplementary Figure 1 -*** *Similarity of cortical reconstructions across different T1w acquisition protocols. For each subject, pre- and postoperative cortical thickness was averaged across the contralateral hemisphere where limited change is expected.*

### SUPPLEMENTARY METHODS - Clinical Workflow

The following sections *(decision of surgical technique, surgical technique of ATL, surgical technique of tsSAHE)* have been published previously elsewhere[1] and are reproduced here for completeness of content.

***Decision of the Surgical Technique***

All patients were discussed in an interdisciplinary epilepsy surgery conference and decision regarding the type of surgery *(transsylvian selective amygdalohippocampectomy [tsSAHE] vs. anterior temporal lobectomy [ATL])* was made based on the results of presurgical evaluation: Patients with unilateral interictal epileptiform discharges, ictal EEG-patterns, and seizure semiology, as well as clinical lateralising signs corresponding to mesial temporal lobe origin and absence of any other pathology in extramesial temporal lobe on MRI, were subjected to tsSAHE; patients without clearly circumscribed interictal temporal spikes (i.e. with broad field or bitemporal distribution), ictal EEG-patterns, and seizure semiology, as well as clinical lateralising signs which were not clearly corresponding to a mesial temporal onset and extramesial temporal lobe structural changes on MRI, were subjected to ATL.

***Surgical Technique of ATL***

The posterior extent of the resection was set at 3-4cm on the dominant *(13 patients)* and 4-5cm on the non-dominant side *(19 patients)* measured along T2 from the pole[2]. The lateral corticotomy was extended to the basal temporal lobe, the ventricle was accessed through the T2/T3/T4-incision. Subpial dissection was performed along T1 to the inferior insular sulcus and down to the uncus. The dorsal amygdala was transected by blunt dissection connecting the anterior choroidal point with the insular limen, respecting the pial barrier below the stem of the middle cerebral artery. By extending the incision through the ependyma along the collateral sulcus, posterior parts of the parahippocampal gyrus and hippocampus were disconnected. The resected specimen usually consisted of a polar and a medial bloc of tissue. A hippocampal-sparing left-sided resection was performed in a single patient.

***Surgical Technique of tsSAHE***

In all patients, the transsylvian-transamygdala route was used[3–5]. After opening the sylvian fissure, a corticotomy on the temporal side of the limen insulae was performed. By resection and suction of the amygdala and adjoining uncal cortex up to the level of the optic tract, the anterior roof of the temporal horn was opened. The posterior margin of the resection was defined by a transverse transependymal cut through the most posterior part of the hippocampus. The collateral sulcus was exposed through the ependyma lateral to the hippocampus. This incision was connected to the posterior transverse hippocampal incision, and the lateral limit of the resection was defined by following the collateral sulcus rostrally, taking into account information on the individual sulcal pattern provided by MRI. After dissecting around the hippocampal head in a subpial plane, cutting the unco-hippocampal arteries passing through the uncal sulcus, and lastly coagulating and cutting the vein usually present at the anterior end of the choroid fissure and draining into the basal vein, the parahippocampal-hippocampal specimen was removed en-bloc. No retractor is used at any time point during the procedure.

### SUPPLEMENTARY METHODS - Baseline characteristics

***Cohort characteristics***

Fifty-nine patients fulfilled the inclusion criteria included in this study (ATL:SAHE=31:28*)*. Overall, no significant differences were found between treatment both groups *(Supplementary Table 1)*, except that more male patients underwent ATL.

| **Characteristics** | **tsSAHE** *(n=28)* | **ATL** *(n=31)* | **p-value** |
|---|---|---|---|
| Sex (male:female) | 12:16 | 22:9 | 0.02[1] |
| Disease duration until surgery (median years, IQR) | 12 (25) | 15 (20) | 0.93[2] |
| Age at disease onset (mean age, SD) | 20 (16) | 19 (12) | 0.84[3] |
| Age at surgery (mean age, SD) | 39 (11) | 38 (12) | 0.70[3] |
| TLE side (left:right) | 16:12 | 11:20 | 0.10[1] |
| MRI diagnosis | | | |
| 1. MRI negative | 2 | 15 | |
| 2. HS | 11 | 3 | |
| 3. HA | 1 | 4 | |
| 4. HS + HA | 10 | 4 | |
| 5. FCD | 1 | 1 | |
| 6. LGG | 1 | 1 | |
| 7. Other | 2 | 3 | |
| Histological diagnosis | | | |
| 1. HS | 23 | 9 | |
| 2. FCD | 1 | 5 | |
| 3. HS + FCD | 1 | 3 | |
| 4. mMCD | 0 | 3 | |
| 5. Ganglioglioma | 0 | 1 | |
| 6. Other | 3 | 9 | |
| 7. Lost to FU | 0 | 1 | |
| Presurgical MRI (median months, IQR) | 7 (7) | 7 (8) | 0.90[2] |
| Postsurgical MRI (median months, IQR) | 4 (1) | 5 (3) | 0.65[2] |

***Supplementary Table 1*** *- Baseline characteristics of the reported cohort. ATL = anterior temporal lobectomy; FCD = focal cortical dysplasia; FU = follow up; HA = hippocampal atrophy; HS = hippocampal sclerosis; IQR = interquartile range; LGG = low grade glioma; mMCD = mild malformation of cortical development; SD = standard deviation; tsSAHE = transsylvian selective amygdalohippocampectomy; Statistical Tests: 1= Qui-squared test, 2 = Mann-Whitney U test, 3 = Student's t-test. Adapted from*[1]

#### SUPPLEMENTARY METHODS – Imaging Pipeline

##### *Image processing*

Processing of diffusion data was performed using MRTrix3[6] (v3.0.3), processing of structural data using Freesurfer[7] (v7.4.1) and co-registrations using ANTs[8] (v2.3.5) unless indicated otherwise. Diffusion data was denoised using Marchenko-Pastur Principal Component Analysis[9] and corrected for Gibbs ringing based on local subvoxel-shifts[10]. Susceptibility distortions and eddy currents were corrected[11,12] using a synthetical b0 image predicted from T1w data, generated with the SyNb0-DiSCo tool[13]. Finally, data was corrected for B1 field inhomogeneities using ANTs N4 bias correction[14]. Tissue response functions were estimated from preoperative data by the *dhollander* algorithm[15] and averaged to group-wise responses. We performed single-shell-three-tissue constrained spherical deconvolution[16] to estimate orientation distribution functions (ODFs) that we intensity normalised across subjects[17]. Structural data were N3 bias field corrected[18], intensity normalised[19] and rigidly registered to preprocessed diffusion images.

##### *Template generation*

For each subject, we generated subject-specific unbiased[20] templates across sessions. Image registration was based on structural data: First, we rigidly aligned sessions using robust template registration[21] as implemented in the longitudinal Freesurfer processing stream. Rigidly aligned structural images were then further refined with ANTS' deformable symmetric normalisation[8]. Next, spatially aligned structural images were then averaged to an unbiased template. Given that parts of the brain were removed in-between sessions, averaging image intensities in areas of tissue resection inevitably results in unplausible image intensities. For these areas, we therefore imputed intensity values from preoperative images, resulting in an anatomically intact structural template across sessions *(see Supplementary Figure 2)*. The previously generated transforms were used to spatially align, reorient[22], and modulate[23] fODF images. Postoperative fODFs images were NaN masked in areas of tissue resection and sessions were averaged to generate fODF templates.

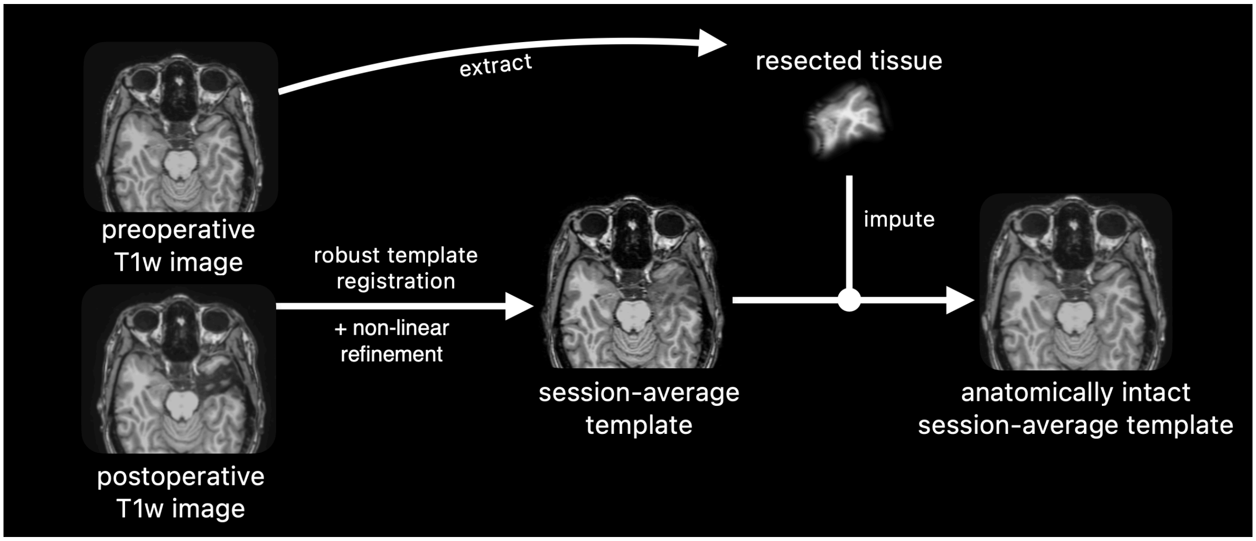


***Supplementary Figure 2*** *– Anatomically intact template generation using structural data. Structural session data is spatially aligned and averaged. In areas of tissue resection preoperative tissue was imputed to avoid unplausible average estimates.*

***Resection cavity segmentation***

The resection cavity was segmented from spatially aligned structural images using the following approach: first, images were segmented into three tissues, namely white matter *(WM)*, grey matter *(GM),* and cerebrospinal fluid *(CSF)* using Atropos' K-Means clustering[24]. After smoothing, we calculated the difference between CSF segmentations and thresholded them at 0.2, effectively discarding voxels with a longitudinal CSF difference probability below 20%. The segmentation was then multiplied by an intersection mask to discard any field-of-view-related differences, followed by erosion to disconnect clusters with single-voxel connections, selection of the largest component, and dilation of the cluster back to its original size. This segmentation was then multiplied by a preoperative parenchyma mask *(combined preoperative WM + GM segmentation)* to only keep voxels that were classified as tissue in the preoperative image. The preliminary mask was then further refined, repeating several above-mentioned steps: within the mask, Atropos' K-Means clustering was performed, separating voxels into two clusters. The result is a sub-segmentation of the purely CSF-filled resection cavity and remaining clusters of tissue not classified as such during the first iteration. After erosion, selection of the largest component, and dilation back to its original size, we proceeded with the purely CSF filled sub-segmentation as surgical defect.

***Further resection cavity refinement***

We found that small registration errors can lead to clusters of stray voxels that are part of the resection but were misclassified as non-resected tissue in the atlas labelling of the unbiased template. To avoid such voxels driving longitudinal cortical thickness estimates, we relabelled them as part of the resection if they were only present in either the preoperative or postoperative spatially aligned tissue segmentation. Furthermore, we relabelled ipsilateral brain regions with voxel cluster sizes <5% of their contralateral counterpart as part of the resection, avoiding that results are driven by small remnants of otherwise resected brain areas.

### SUPPLEMENTARY METHODS – Adaption of the longitudinal Freesurfer pipeline for surgical image data

***Processing of longitudinal structural data***

We adapted Freesurfer's longitudinal processing stream[20,21,25] according to Hoffman et. al[26], who showed that in the presence of large-scale anatomical changes, it is preferable to estimate surfaces from a non-linear base template.  We therefore initialised the longitudinal processing stream and injected the previously generated *(non-linearly refined and resection-corrected)* templates, allowing the reconstruction of anatomically intact session-average timepoints. Note that while a non-linear template is used for anatomical reconstruction, the longitudinal processing stream still uses the initially generated rigid transforms to transfer information between base template and timepoints; this methodology is coherent with the processing stream used in Hoffman et. al. and is based on their observation that use of non-linear transforms can be detrimental due to non-ideal alignment around the cortex. Since the base template's surfaces only serve as a starting point for further refinement to fit surfaces to session data, remaining misalignments arising from the use of rigid transformations will be corrected by adjusting vertex positions to align with the respective time point data. Finally, to avoid extraction of measures from resected regions, we segmented Desikan-Killiany atlas labels into resected and non-resected sub-parcels.

### SUPPLEMENTARY METHODS – SIFT2$_{unbiased}$: a novel framework for unbiased white matter density mapping in individuals

***Processing of longitudinal diffusion data***

The reconstruction of structural connectomes is complex and involves iterative methods that are highly sensitive to individual starting points and noise present in the data. This results in significant methodological variance when cross-sectionally reconstructing longitudinal timepoints[27], making it impossible to distinguish method variance from biological effects. For the estimation of longitudinal metrics from T1w data, it has proven beneficial to avoid independent reconstruction steps by estimating the average gross anatomy, which is then further refined on session-specific data[25]. Translating this concept to dMRI analysis, we here introduce SIFT2$_{unbiased}$ *(SIFT=Spherical Deconvolution Based Filtering of Tractograms),* an unbiased framework to longitudinally estimate white matter density weights in individuals. The framework is an extension of the commonly used SIFT2 algorithm[28] and avoids independent reconstruction of multiple timepoints by reconstructing an weighted base streamline tractogram within an unbiased template, which is then refined to fit session-specific fibre densities *(see Supplementary Fig. 3).*

For this study, we constructed SIFT2 weighted tractograms of 10 million streamlines from an unbiased template across sessions using dynamic seeding[28] and anatomically constrained tractography[29] *(ACT).* This weighted tractogram was then used to initialise SIFT2 tractogram optimisation of spatially aligned, reoriented, and fODF modulated pre- and postoperative timepoints. Before initialising postoperative optimisation, we additionally set weights of streamlines intersecting the resection mask to zero, effectively excluding surgically resected connections from optimisation. Finally, per-streamline white matter density weights were summarised to node-wise structural connectomes to assess longitudinal differences in Fibre Bundle Capacity[30] (FBC). Note that features obtained from structural data within this connectome pipeline, namely ACT's five-tissue-types image as well as the atlas segmentation, were also obtained from an unbiased structural template within the same space[20], further reducing independent reconstruction steps.

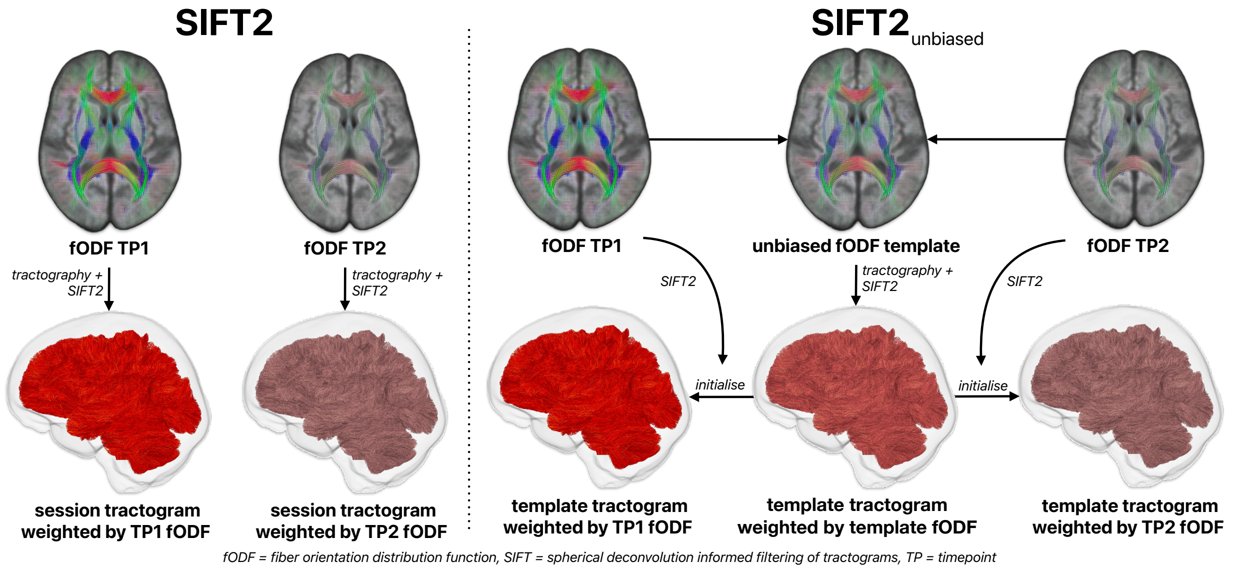


***Supplementary Figure 3*** *– Overview SIFT2 vs. SIFT2$_{unbiased}$ for longitudinal quantification of per-streamline white matter densities. Left: Independent reconstruction of SIFT2 weighted tractograms. Right: Unbiased reconstruction using a weighted base tractogram to initialise SIFT2 optimisation of timepoints. fODF=fibre orientation distribution function, TP=timepoint, SIFT=spherical deconvolution informed filtering of tractograms.*

#### SUPPLEMENTARY RESULTS – SIFT2 vs. SIFT2$_{\text{unbiased}}$

Unbiased connectome reconstruction delineated longitudinal decreases in FBC that were most pronounced in the hemisphere ipsilateral to the resection, as would be biologically expected. Independent reconstruction on the other hand led to dubious increase and decreases within both hemispheres. We showed this to be the result of unaccounted reconstruction variance *(see Extended Data Fig. 3).* To further demonstrate this point, we projected log-transformed pre- and postoperative connectomes of the contralateral hemispheres *(where limited biological change is expected)* into a common two-dimensional UMAP space *(metric: correlation, settings: default)*, where timepoints that are more similar to each other map closer together. The analysis confirmed greater similarity of timepoints estimated using the unbiased framework *(Euclidean distance 0.14±0.07)* as compared to those estimated using cross-sectional reconstruction *(Euclidean distance 0.31±0.34, see Supplementary Fig. 4).*

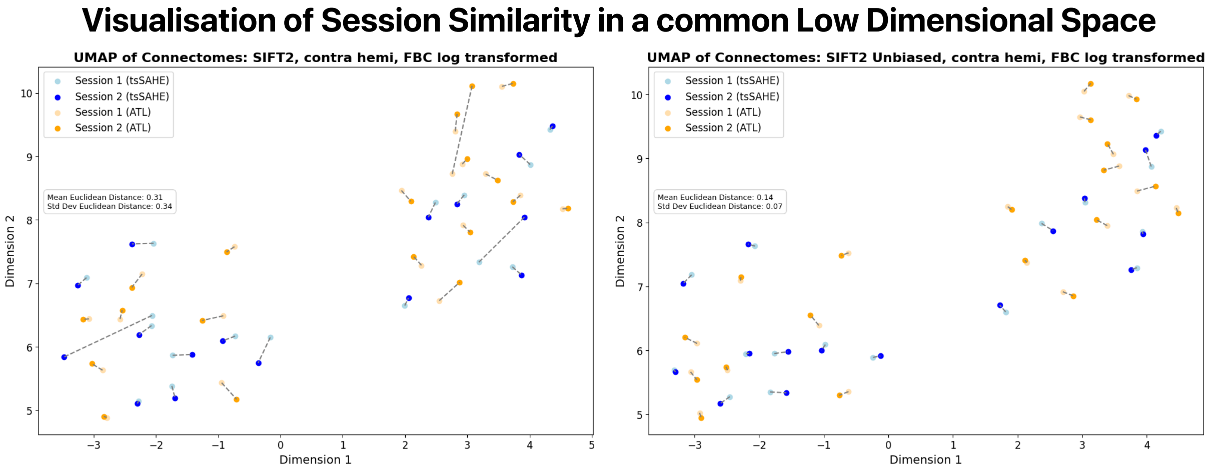


***Supplementary Figure 4*** *– Projection of contralateral connectome data into a common low-dimensional space. Unbiased reconstruction led to greater similarity between timepoints as compared to independent reconstruction.*

1. Pruckner, P. *et al.* Visual outcomes after anterior temporal lobectomy and transsylvian selective amygdalohippocampectomy: A quantitative comparison of clinical and diffusion data. *Epilepsia* **64**, 705–717 (2023).
2. Zentner, J. *Surgical Treatment of Epilepsies*. (Springer, 2002).
3. Yaşargil, M. G., Türe, U. & Yaşargil, D. C. Impact of temporal lobe surgery. *Journal of neurosurgery* **101**, 725–738 (2004).
4. Dorfer, C. *et al.* Mesial temporal lobe epilepsy: long-term seizure outcome of patients primarily treated with transsylvian selective amygdalohippocampectomy. *Journal of neurosurgery* **129**, 174–181 (2017).
5. Yasargil, M., Teddy, P. & Roth, P. Selective amygdalo-hippocampectomy operative anatomy and surgical technique. *Advances and Technical Standards in Neurosurgery: Volume 2* 93–123 (1985).
6. Tournier, J.-D. *et al.* MRtrix3: A fast, flexible and open software framework for medical image processing and visualisation. *NeuroImage* **202**, 116137 (2019).
7. Fischl, B. FreeSurfer. *NeuroImage* **62**, 774–781 (2012).
8. Avants, B. B. *et al.* A reproducible evaluation of ANTs similarity metric performance in brain image registration. *NeuroImage* **54**, 2033–2044 (2011).
9. Veraart, J. *et al.* Denoising of diffusion MRI using random matrix theory. *NeuroImage* **142**, 394–406 (2016).
10. Kellner, E., Dhital, B., Kiselev, V. G. & Reisert, M. Gibbs-ringing artifact removal based on local subvoxel-shifts. *Magnetic Resonance in Medicine* **76**, 1574–1581 (2016).
11. Smith, S. M. *et al.* Advances in functional and structural MR image analysis and implementation as FSL. *NeuroImage* **23**, S208–S219 (2004).
12. Andersson, J. L. R. & Sotiropoulos, S. N. An integrated approach to correction for off-resonance effects and subject movement in diffusion MR imaging. *NeuroImage* **125**, 1063–1078 (2016).
13. Schilling, K. G. *et al.* Synthesized b0 for diffusion distortion correction (Synb0-DisCo). *Magnetic Resonance Imaging* **64**, 62–70 (2019).
14. N. J. Tustison *et al.* N4ITK: Improved N3 Bias Correction. *IEEE Transactions on Medical Imaging* **29**, 1310–1320 (2010).

15. Dhollander, T., Raffelt, D. & Connelly, A. *Unsupervised 3-Tissue Response Function Estimation from Single-Shell or Multi-Shell Diffusion MR Data without a Co-Registered T1 Image*. (2016).
16. Dhollander, T., Mito, R., Raffelt, D. & Connelly, A. *Improved White Matter Response Function Estimation for 3-Tissue Constrained Spherical Deconvolution*. (2019).
17. Raffelt, D. *et al. Bias Field Correction and Intensity Normalisation for Quantitative Analysis of Apparent Fibre Density*. (2017).
18. J. G. Sled, A. P. Zijdenbos, & A. C. Evans. A nonparametric method for automatic correction of intensity nonuniformity in MRI data. *IEEE Transactions on Medical Imaging* **17**, 87–97 (1998).
19. Dale, A. M., Fischl, B. & Sereno, M. I. Cortical Surface-Based Analysis: I. Segmentation and Surface Reconstruction. *NeuroImage* **9**, 179–194 (1999).
20. Reuter, M. & Fischl, B. Avoiding asymmetry-induced bias in longitudinal image processing. *NeuroImage* **57**, 19–21 (2011).
21. Reuter, M., Rosas, H. D. & Fischl, B. Highly accurate inverse consistent registration: A robust approach. *NeuroImage* **53**, 1181–1196 (2010).
22. Raffelt, D., Tournier, J.-D., Crozier, S., Connelly, A. & Salvado, O. Reorientation of fiber orientation distributions using apodized point spread functions. *Magnetic Resonance in Medicine* **67**, 844–855 (2012).
23. Raffelt, D. *et al.* Apparent Fibre Density: A novel measure for the analysis of diffusion-weighted magnetic resonance images. *NeuroImage* **59**, 3976–3994 (2012).
24. Avants, B. B., Tustison, N. J., Wu, J., Cook, P. A. & Gee, J. C. An Open Source Multivariate Framework for n-Tissue Segmentation with Evaluation on Public Data. *Neuroinformatics* **9**, 381–400 (2011).
25. Reuter, M., Schmansky, N. J., Rosas, H. D. & Fischl, B. Within-subject template estimation for unbiased longitudinal image analysis. *NeuroImage* **61**, 1402–1418 (2012).
26. Hoffmann, M., Salat, D., Reuter, M. & Fischl, B. Longitudinal FreeSurfer with non-linear subject-specific template improves sensitivity to cortical thinning. *Athinoula A. Martinos Center for Biomedical Imaging, Charlestown, MA, United States; Department of Radiology, Harvard Medical School, Boston, MA, United States; German Center for Neurodegenerative Diseases, Bonn, Germany; Computer Science and Artificial Intelligence Laboratory, Massachusetts Institute of Technology, Cambridge, MA, United States*.

27. Smith, R. E., Tournier, J.-D., Calamante, F. & Connelly, A. The effects of SIFT on the reproducibility and biological accuracy of the structural connectome. *NeuroImage* **104**, 253–265 (2015).
28. Smith, R. E., Tournier, J.-D., Calamante, F. & Connelly, A. SIFT2: Enabling dense quantitative assessment of brain white matter connectivity using streamlines tractography. *NeuroImage* **119**, 338–351 (2015).
29. Smith, R. E., Tournier, J.-D., Calamante, F. & Connelly, A. Anatomically-constrained tractography: Improved diffusion MRI streamlines tractography through effective use of anatomical information. *NeuroImage* **62**, 1924–1938 (2012).
30. Smith, R. E., Raffelt, D., Tournier, J.-D. & Connelly, A. Quantitative streamlines tractography: methods and inter-subject normalisation. *Aperture Neuro* 1–25 (2022).
